# How density dependence, genetic erosion, and the extinction vortex impact evolutionary rescue

**DOI:** 10.1101/2022.10.29.514126

**Authors:** Scott W. Nordstrom, Ruth A. Hufbauer, Laure Olazcuaga, Lily F. Durkee, Brett A. Melbourne

## Abstract

Following severe environmental change that reduces mean population fitness below replacement, populations must adapt to avoid eventual extinction, a process called evolutionary rescue. Models of evolutionary rescue demonstrate that initial size, genetic variation, and degree of maladaptation influence population fates. However, many models feature populations that grow without negative density dependence or with constant genetic diversity despite precipitous population decline, assumptions likely to be violated in conservation settings. We examined the simultaneous influences of density-dependent growth and erosion of genetic diversity on populations adapting to novel environmental change using stochastic, individual-based simulations. Density dependence decreased the probability of rescue and increased the probability of extinction, especially in large and initially well-adapted populations that previously have been predicted to be at low risk. Increased extinction occurred shortly following environmental change, as populations under density dependence experienced more rapid decline and reached smaller sizes. Populations that experienced evolutionary rescue lost genetic diversity through drift and adaptation, particularly under density dependence.

Populations that declined to extinction entered an extinction vortex, where small size increased drift, loss of genetic diversity, and the fixation of maladaptive alleles, hindered adaptation, and kept populations at small densities where they were vulnerable to extinction *via* demographic stochasticity.

## Introduction

Adaptation to novel environments is often necessary for populations to persist in this current era of large-scale anthropogenic change and habitat alteration. Novel or sudden environmental change can abruptly render a population poorly adapted to its habitat. When the environmental change is so severe that mean population fitness falls below the replacement rate, the population will go extinct if it does not adapt. This process of populations adapting to severe environmental change sufficiently to avoid extinction is called evolutionary rescue [1,2].

Gomulkiewicz and Holt [1] formalized the idea of evolutionary rescue, highlighting the “U”– shaped demographic trajectory where populations initially decline due to maladaptation to novel environmental conditions, followed by adaptation that allows populations to avoid extinction and return to their original size. They found that evolutionary rescue was most likely to occur in large populations with high genetic diversity and a relatively low degree of maladaptation to their environment (i.e., population mean phenotype close to that favored by the novel environment) at the time of the environmental change. Subsequent work in controlled experiments has confirmed the importance of these factors [3–5] to evolutionary rescue. However, these models and laboratory results often do not include features of population regulation or adaptive processes present in wild populations, hindering their predictive ability in conservation settings [2,6].

Models examining rescue often rely on assumptions of density-independent growth [7], potentially making the findings less applicable to many conservation targets. For example, Gomulkiewicz and Holt’s model [1] assumes that population growth is density independent and that populations can grow without constraint. This assumption will often be violated in conservation settings where habitat loss, degradation, or other environmental change limits resource availability and increases intraspecific competition. For example, a recent analysis of 73 populations of rare plant species revealed that negative density dependence was statistically detectable in a majority of them, emphasizing the importance of considering it in population projection models [8].

Additionally, models often assume that genetic variance in the population remains constant. This is present in several models (e.g., [1,9]) relying on quantitative genetics results from Lande [10], which assume constant genetic variance. However, threatened populations experiencing rapid decline are likely to experience erosion of genetic diversity [11–13] that may decrease their ability to adapt to novel conditions [14], possibly hindering rescue. Furthermore, independently of drift or other sources of stochasticity, populations experiencing directional or stabilizing selection due to environmental change will lose genetic variation as they adapt [15]. This loss of variation reduces the “variance load”, which can be defined as reduced population growth due to the mean distance of individual phenotypes from the optimum [16]. Thus, reductions in genetic variance will be the norm in small populations experiencing novel selection pressures and can be associated with either decreases in fitness due to drift load or increases in fitness due to adaptation and reduced variance load, limiting or improving the potential for evolutionary rescue, respectively.

Simultaneously incorporating realistic demographic and genetic processes such as negative density dependence and genetic erosion will improve the accuracy of predictions from models of evolutionary rescue for populations and species of conservation concern. Negative density dependence that is caused by intraspecific competition should constrain evolutionary rescue by accelerating population declines and slowing population recovery from reduced size [9]. Some models of rescue and environmental tracking have included constrained population size by including a ceiling-type carrying capacity (e.g., [17–20]), but growth below carrying capacity in these models was density independent, failing to capture how intraspecific competition or other density-dependent processes depress population growth or influence genetic diversity in declining and recovery phases of rescue. Orr and Unckless [21] briefly analyzed a density-dependent model of adaptation with a single locus trait, finding that density dependence impedes rescue by reducing the survival of novel beneficial mutants. However, they did not investigate effects of density dependence on adaptation from standing genetic variation. This ignores the influence of drift, which at reduced population size can cause the loss of favored alleles (but see [22] for possible increase in frequency). Chevin and Lande [9] found that density dependence impedes rescue by accelerating population decline. However, their model held genetic diversity constant, ignoring drift and potentially mischaracterizing the rate at which populations adapt and return to their pre-change density. A population’s additive genetic variance for a trait influences the trait’s rate of adaptation under directional selection [10,14]. Loss of this genetic diversity as populations decline in response to environmental change should slow rates of adaptation, increasing the time until rescue or making eventual rescue less likely. While some analytical models incorporate effects of drift due to finite population size, they do so in a manner that only produces variance around the mean rate of adaptation (e.g., [18]), still holding genetic and phenotypic variance constant and leaving the expected rate of adaptation unaffected. Some rescue models allow genetic diversity to change over time, finding that drift could potentially slow adaptation [23,24]. But, these models do not include interactions of changing genetic diversity with competition that constrains population growth.

Modeling these demographic and genetic processes simultaneously is important, as they can interact and reinforce each other, creating a positive feedback loop of demographic and genetic decline. In the conservation biology literature this positive feedback loop is called an “extinction vortex” [25]. If populations in the process of evolutionary rescue are drawn into such loops, adaptation and rescue will be impeded. In an extinction vortex (Fig. 1) small populations experience drift and inbreeding, reducing fitness directly *via* increased genetic load and indirectly by erosion of genetic variation and reduced rates of future adaptation [11], keeping populations small or causing them to decline further to a smaller size. This in turn leads to further genetic erosion or increased genetic load, reduced fitness, and further declines in population size until inevitably the population is highly vulnerable to extinction *via* stochasticity in births and deaths, demographic heterogeneity, and sampling variation in sex ratios [26]. This feedback loop potentially opposes or resists the evolutionary rescue process, where adaptation increases fitness, increasing population growth rates to the point where population size stabilizes then eventually returns to pre-change levels. Negative density dependence has the potential to disrupt evolutionary rescue and draw populations back towards the vortex by accelerating reductions in, or slowing increases in, population growth rate, fitness, and genetic diversity (Fig. 1). The original conceptualization of an extinction vortex does not necessarily assume a degraded or altered habitat where adaptation is necessary for persistence [25].

**Figure 1:**
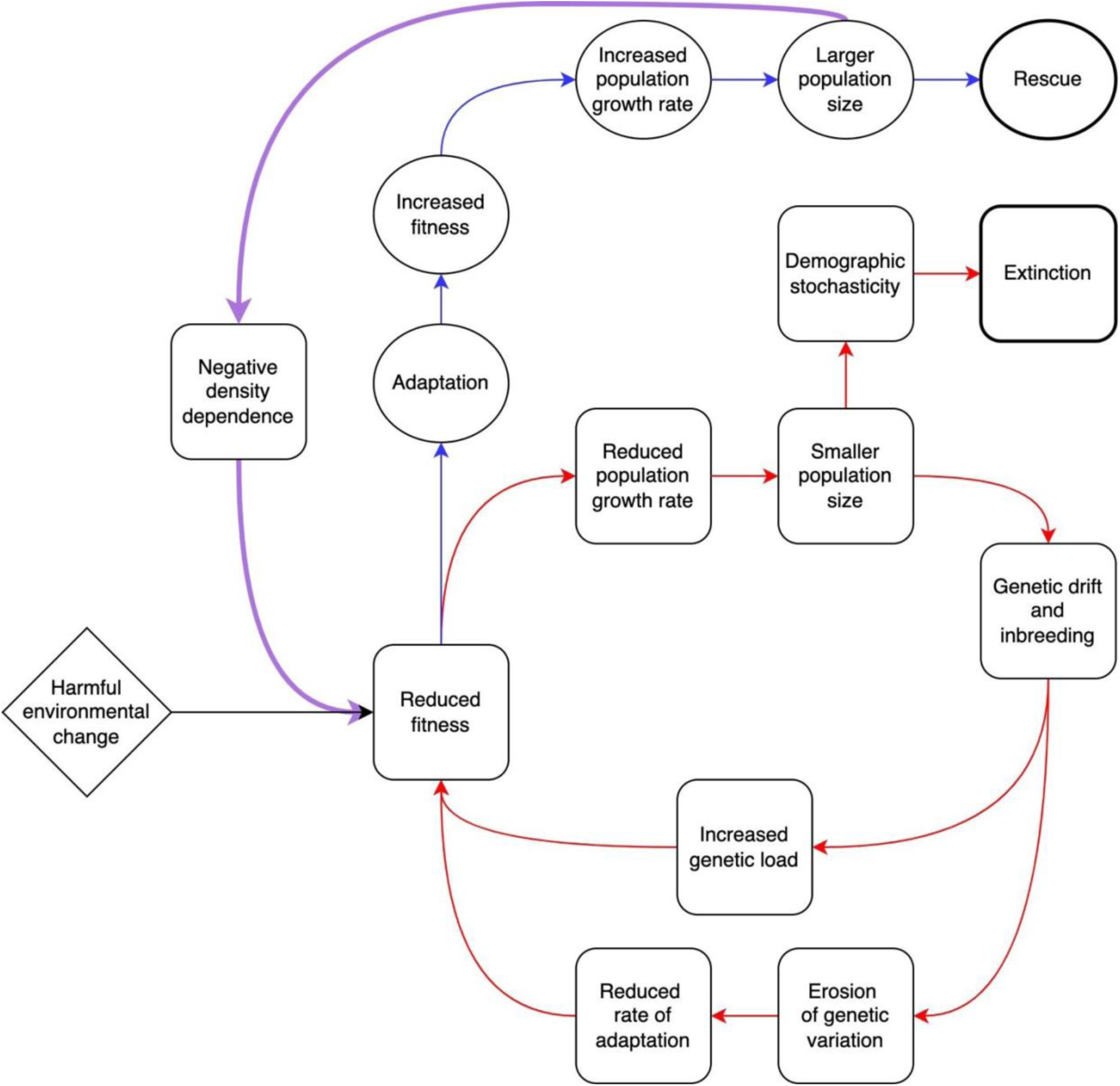
Event sequence diagram of evolutionary rescue (blue, ovals) and two connected extinction vortices (red, squares) that populations may face following harmful environmental change. The effects of negative density dependence, potentially impeding rescue and drawing populations into a vortex, are indicated in purple.

Perhaps for this reason, to our knowledge the extinction vortex has not been explicitly studied in the context of evolutionary rescue and adaptation to environmental shifts. Yet, these interactive, combined effects of small population size could be of particular importance in evolutionary rescue, as by definition populations undergoing rescue initially decline to smaller sizes, which could initiate entry into an extinction vortex. The extinction vortex also necessitates understanding how density dependence and genetic erosion interact with other important determinants of the success of rescue, e.g., the initial conditions with respect to population size, genetic diversity, and degree of maladaptation [1,3,4]. For example, large populations adapting to sudden environmental change might rarely approach the extinction vortex under density-independent growth but might more often enter the vortex under density dependence. Ultimately, under realistic conditions in natural settings, evolutionary rescue might often involve a population escaping from an extinction vortex.

Here, we relax the assumptions of density independence and constant genetic diversity to examine their simultaneous effects on evolutionary rescue and the extinction vortex, establishing a connection between these two major concepts in conservation biology. We do this by combining stochastic Ricker population growth and a genetic framework with a finite-locus quantitative trait in the same individual-based model. We simulate both rapid (15 generations) and longer (50 generations) timescales. We address two main objectives. First, we assess how negative density dependence interacts with initial conditions with respect to population size, additive genetic variance, and degree of genetic maladaptation in the novel environment to influence trajectories of population size, probabilities of extinction and rescue, additive genetic variance, and fitness over time. Second, we test for the presence of an extinction vortex and evaluate how negative density dependence influences this vortex.

## Methods

### Model description

We explored evolutionary rescue and extinction using stochastic simulations. This approach allowed us to incorporate several stochastic demographic and genetic processes that are important for rescue or extinction [2,26]. We derive a discrete-time individual-based model with diploid genetic inheritance that has general features of the semelparous life history strategy common to many sexually reproducing organisms. We use non-overlapping generations for comparison with other models of adaptation and evolutionary rescue for semelparous organisms (e.g., [1,9,23]). Our genetic submodel is a finite quantitative locus model with random and independent segregation of alleles, similar to that of Boulding and Hay [23]. Our demographic submodel is based on an expanded stochastic Ricker model [26,27], which models density dependence.

Each individual has an intrinsic fitness determined by genetic and environmental components. The genetic component is determined by each of *m* biallelic diploid loci. Each allele-copy contributes a positive 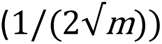 or negative 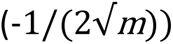 amount to the genotype of individual *i*, *g_i_*, which is the sum of allele-copies across all loci. The additive genetic variance in the population is thus

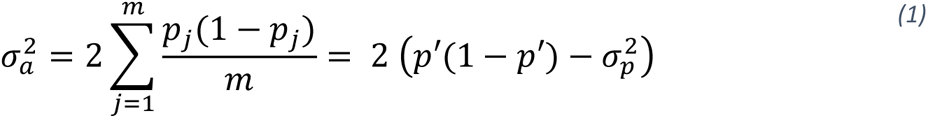

where *p_j_* is the frequency of the positive allele at locus *j* across the population and ρ′ and 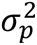 are respectively the mean and variance of allele frequencies across all loci (see [28] and Supporting Information A). We do not model mutations, noting that in our simulated experimental design (see below) populations are small and timespans are short, such that at realistic mutation rates mutation would be rare in our simulations. An individual’s phenotype, *z_i_*, is the genotype, *g_i_*, plus a deviation due to environmental variation drawn from a normal distribution with mean zero and variance 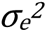. Intrinsic fitness, *W_i_*, the expected fitness of an individual in the absence of density effects, is determined by Gaussian selection [10] such that *W_i_* = *W*_*max*_ *exp*(−(*Z*_*i*_ − θ)^2^/2*w*^2^), where θ is the optimal phenotype, *w^2^* (*w*) is the variance (width) of the fitness landscape and is inversely proportional to selection strength, and *W_max_* is the maximum intrinsic fitness (i.e., intrinsic fitness of an individual with the optimal phenotype). The expected number of offspring of an individual, *R_i_*, is determined according to a Ricker function that reduces fitness due to density dependence such that *R*_*i*_ = *W_i_e*^−αN^, where *α* is the strength of density dependence and *N* is the population size [27]. Every generation, each female mates with one male chosen uniformly at random from the population. Each female produces a Poisson-distributed number of offspring with mean 2*R_i_*, with parental alleles segregated independently and at random to offspring. Sex is assigned to each individual upon birth with equal probability. Generations are not overlapping, i.e., only one generation is present in each time step. As such, if all females have zero offspring, or if all individuals in the population are of the same sex, then extinction is guaranteed in the next time step.

### Experimental Design, Simulations, and Analysis

While extinction is straightforward to identify, the classical criterion of evolutionary rescue (population growth rate above replacement) is more difficult to operationalize for individual populations because rescued populations may transiently decline in size or growth rate due to demographic or genetic stochasticity and revert to an “unrescued” state. We operationalize rescue using two separate criteria: (1) expected number of offspring per individual (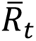, as defined below) greater than 1 for three consecutive generations (hereafter “fitness-based rescue”) and (2) populations exceeding their initial size for three consecutive generations (hereafter “size-based rescue”).

Parameter values are given in Supporting Information Table B1, justified in full in Supporting Information C, and are realistic for a variety of natural populations [29,30]. We simulated populations with *m* = 25 loci and an environmental change that moved the phenotypic optimum from its prior state of 0 to θ = 2.8. Because θ > 0, the environment in our simulations favors a shift toward positive alleles. Populations were founded with positive and negative alleles assigned to each individual’s loci with equal probability such that across simulation trials the expected population mean genotype was zero, although due to sampling there was variation in initial population mean genotype across trials. Thus, to reach the new phenotypic optimum θ = 2.8, a typical population needed to change from 25 positive allele copies (out of 50) per individual to 39 allele copies per individual (see Supporting Information D). All simulations were run with maximum intrinsic fitness *W_max_* = 2, non-genetic phenotypic variance 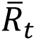 = 0.5, and *w^2^* = 3.5 as the variance of the fitness landscape. With these parameter values, mean intrinsic fitness in the typical population immediately following the environmental change was approximately 0.65 such that populations were sufficiently challenged to the point that both rescue and extinction were possible on rapid timescales.

To evaluate the effects of negative density dependence on evolutionary rescue and its potential interactions with initial population size and genetic diversity, we conducted a multi-factorial simulation experiment (2 x 2 x 2, i.e., eight treatments). Populations were simulated with the following parameter values: negative density dependence absent (*α* = 0) or present (*α* = 0.0035, carrying capacity approx. 200), hereafter referred to as “density-independent populations” or “density-dependent populations”; initial size *N*_0_ = 100 or *N*_0_ = 20, hereafter referred to as “large” or “small” populations in reference to their initial state rather than their eventual size; initial genetic diversity high (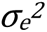 ≈ 0.5, heritability approx. 0.5) or low (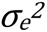≈ 0.25, heritability approx. 0.33 [28]), hereafter “high diversity” or “low diversity” populations, again in reference to their initial state rather than their eventual level of diversity. These initial heritabilities match the range of heritabilities in life history traits in wild populations [31]. Note that large populations do not necessarily harbor more genetic diversity than small populations independently of the genetic diversity treatment. To simulate populations with initially low diversity, at the beginning of each simulation six loci were set to fixation for the positive (adaptive) allele and six were set to fixation for the negative (maladaptive) allele, leaving only 13 loci (approximately half of the genome) available for selection to act upon while leaving the expected mean population genotype unchanged compared to the high-diversity treatments. Stochasticity in the founding of populations produced sampling variation in the initial mean population genotype; we took advantage of this sampling variation to also analyze a third important predictor of population fate, initial degree of maladaptation [1,24], and its interactions with density dependence as described in paragraphs below. Due to sampling variation in initial allele frequencies, there was more variation in initial mean population genotypes and phenotypes among small populations than among large populations.

To assess adaptation and extinction on short time scales, we ran 4000 simulations lasting fifteen generations for each treatment. All simulations were performed in R, version 4.1.2 [32]. In each simulation, for each generation we recorded population size (*N_t_*), mean genotype 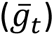 and phenotype 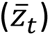, mean intrinsic fitness 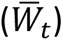, additive genetic variance 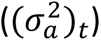, and the proportion of loci at fixation within the population for either allele. Throughout our analysis, we will use an overbar to represent a mean across individuals within one population in one timestep. We define adaptation as an increase in 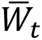. Because fitness is optimized at a positive genotypic value (i.e., θ > 0) and fitness is a monotonic function of maladaptive genetic load, adaptation can be equivalently defined as an increase in 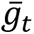 or a reduction in genetic load, 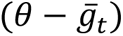.

As a first evaluation of how density dependence, population size, and genetic diversity influence rescue and extinction over 15 generations, we visualized mean population size over time for each treatment. This visualization shows whether the U-shaped population trajectory characteristic of rescue [1] occurs, and if so, how it differs by treatment. All mean population sizes were estimated including extinct populations as size zero, thus incorporating the possibility of extinction into expected population size in each time step. To assess the effects of genetic erosion on population size and extinction, we compared mean population size from each simulated treatment to numerically estimated population size predicted by Gomulkiewicz and Holt’s model [1] (in which phenotypic variance and rate of adaptation are held constant), modified to include a Ricker-density dependence term:

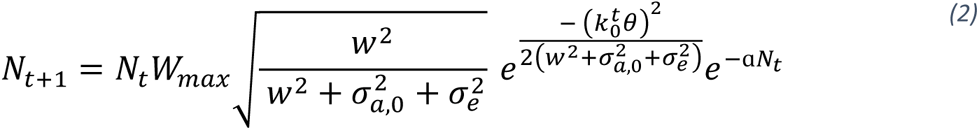

where *k_0_*is the rate of phenotypic change (as described in [1]) in the first time step, averaged across all simulation trials within a genetic diversity treatment.

To quantify the average dynamics of the model we also estimated and visualized the following response variables, averaged over the ensemble of all populations within a treatment: mean population genotype, 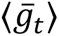, mean intrinsic fitness, 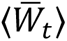, mean additive genetic variance, 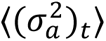, and number of loci at fixation for positive and negative alleles across all populations in each generation. To differentiate ensemble means from population means for a single trial, we use angle brackets (⟨⟩) to denote means of population-level means across an ensemble of simulations. Because these variables are undefined for extinct populations (which have no mean genotype, fitness, or additive genetic variance), estimating unconditional expectations for each generation was impossible, and thus we estimated these means conditioned on survival or extinction at the end of the simulation. Differences in ensemble means for these variables between density-independent and density-dependent populations may be due directly to density dependence, but they may also be due indirectly to non-random extinctions. For example, a population with low mean intrinsic fitness may persist under density independence but go extinct under density dependence, lowering the ensemble mean population fitness for extant populations in a density-independent treatment but not affecting the ensemble mean of extant populations in a density-dependent treatment, confounding comparisons between the treatments. As such, we only use plots of variables conditioned on survival/extinction for qualitative assessment, noting general patterns and interpreting them with caution.

To assess the probabilities of extinction and rescue on longer timescales, we ran 1000 simulations per treatment lasting 50 generations. Due to the computational intensity of simulating large populations, we stopped a trial upon population size exceeding 10000. Because of this censoring, we did not analyze trajectories of demographic or genotypic state variables in these longer simulations, instead focusing only on the probabilities of extinction and rescue. We evaluated extinction in each generation in two ways: (1) instantaneous probability of extinction (the probability of extinction in generation *t* conditioned on surviving to *t*) and (2) cumulative probability of extinction over time (the probability of extinction in or prior to *t*). We also used two separate criteria of rescue (defined above): (1) fitness-based rescue and (2) size-based rescue. We also estimated time to rescue across treatments, defining time to rescue as the first of the three generations where the criterion was met (e.g., a population exceeding original size in generations 10-12 would be considered rescued in generation 10).

We estimated treatment effect sizes on the probabilities of extinction and rescue in the 50-generation trials using Bayesian generalized linear models (GLMs) in the R package *rstanarm,* 2.21.1 [33]. We fit separate models for estimating probabilities of extinction and both types of rescue, treating each simulated population as a Bernoulli response and each of our three treatments (density dependence, initial size, genetic diversity) as categorical variables in each model. Additionally, we estimated effects of initial degree of maladaptation, an important determinant of extinction risk [1,24] by including the mean population genetic load in the initial generation (θ − ̅_0_, henceforth, “maladaptation”) as a continuous explanatory variable. This was possible for our stochastic model because sampling variation in the founding of populations produced random among-trial variation in mean initial genotype. We estimated effects of negative density dependence and its interactions with other variables by fitting a model with a four-way interaction and examining effect sizes, estimated as described in Supporting Information E. The uncertainty around these estimates was sufficiently low and could be arbitrarily improved by running more simulations. An effect size for a categorical variable (for example, initial population size) gives the effect of changing that variable’s parameter on the log odds of extinction (or rescue) while holding all other parameters constant, while an effect size for maladaptation gives the change in the log odds of extinction (rescue) associated with switching five alleles from the positive allele to the negative allele in all individuals.

To test for the presence of an extinction vortex we first reversed and re-centered time for each extinct population to give *τ*, the time until extinction, such that populations had a common origin (*τ* = 0) with consistent meaning across populations [34]. We define *τ* such that *τ* = 0 is the first time step where the population is extinct (i.e., population size of 0 or 1), *τ* = 1 is the last time step in which the population size is above 0 or 1, etc. The extinction vortex is characterized by accelerating declines in population size, genetic variation, and/or rate of adaptation as τ approaches zero [25,34]. Specifically, we calculated the rate of population growth 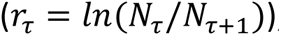, the proportional loss of genetic diversity 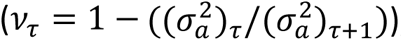, and the rate of adaptation (1 − κ_τ_). κ_τ_ is defined as in [1] such that 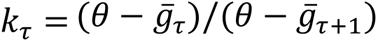; as 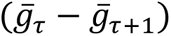 approaches zero (i.e., as the rate of genotypic change approaches zero), κ_τ_ approaches 1 and 1 − κ_τ_ approaches 0. If adaptation proceeds at a constant rate, then κ_τ_ will be constant over time. We used these normalized rates of change instead of non-normalized state variables (e.g., *r*_τ_ instead of *N*_τ_) to facilitate comparisons across treatments and to remove potential biases due to time until extinction. We estimated the ensemble means ⟨*r*_τ_⟩, ⟨ν_τ_⟩, and 1 − ⟨κ_τ_⟩ for all extinct populations in each treatment using the 15-generation simulations, where all state variables were tracked for each population.

Following [34], we considered decreasing ⟨*r*_τ_⟩ (accelerating decline in population size), increasing ⟨ν_τ_⟩ (accelerating loss of genetic diversity), or decreasing (1 − ⟨κ_τ_⟩) (slowing rate of adaptation) in the lead up to extinction (i.e., as *τ* approaches zero) to be evidence of an extinction vortex. Likewise, we assessed how the shape of the extinction vortex varies between density-dependent and density-independent populations qualitatively by noting differences in the mean trajectories of the respective experimental treatments. We did not estimate rates for *τ* = 0 because *r* = –∞ for a population at size zero and because additive genetic variance and mean genotype are undefined for extinct populations.

## Results

### Population size

For all combinations of size and genetic diversity, in each generation density-dependent populations were smaller on average than corresponding density-independent populations (compare black and purple lines, Fig. 2). The minimum average size (bottom of the “U”) reached by density-dependent populations was smaller than that of density-independent populations across treatments, with larger effects in large populations (two-thirds reductions in minimum size) than small populations (one-third reduction). Density-dependent populations were also smaller on average at the end of 15 generations; populations with low diversity had slightly larger reductions in final size (92-95% reduction) than populations with high diversity (85-90% reduction). Regardless of starting population size, density-dependent populations declined for longer and took longer to recover their original size (dotted lines, Fig. 2), if at all.

**Figure 2:**
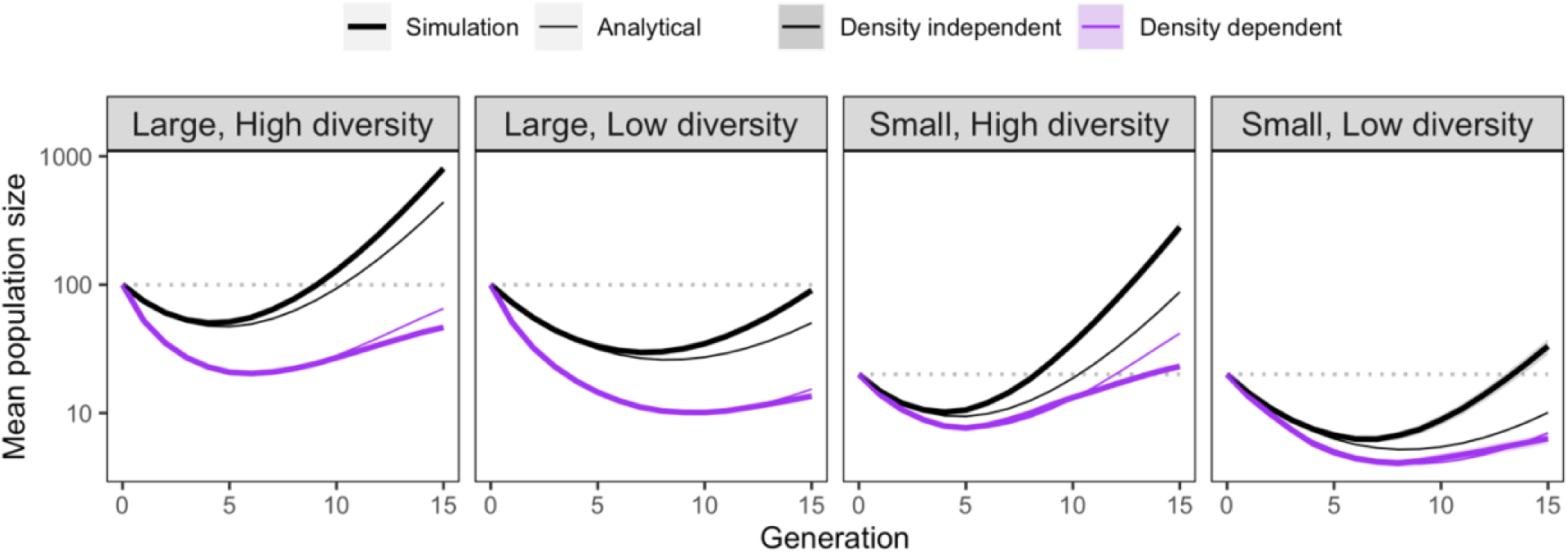
Thick lines show mean population size in each generation for simulated populations (n = 4000 per treatment), including extinct populations as zeros. Thin lines represent expected population size with constant genetic diversity (see equation 2). Shaded areas for the simulation trials are twice the standard error on each side of the mean.

We observed an interaction between density dependence and changing genetic diversity on population size over time. Under density dependence, simulated populations (with changing additive genetic variance) were smaller compared to a model with constant additive genetic variance by the end of simulations (compare bolded and unbolded purple lines, Fig. 2). However, the opposite result was observed for density-independent populations: populations with changing genetic variance were larger than with constant genetic variance (compare bolded and unbolded black lines, Fig. 2). These effects of changing genetic variance emerged for density-independent populations after populations reached their minimum size but took longer to emerge in density-dependent populations. The combined effect was that the effect of density dependence on mean size after 15 generations was larger with genetic erosion than under constant genetic diversity.

### Extinction and rescue

These differences in population size and growth rates over 15 generations translated into more frequent extinctions for populations with density-dependent growth over the course of the 50-generation simulations (Fig. 3). For both density-dependent and density-independent populations, most extinctions occurred in the first 20 generations (Fig. 3*A*,*B*). Density dependence increased extinction probabilities primarily in large populations (Supporting Information Table F1): among large populations, density dependence increased the cumulative odds of extinction 5.0-fold for high-diversity populations and 3.6-fold for low-diversity populations (Fig. 3*Ci*,*ii,* exponentiated effect sizes, Table F1). Among small populations, density dependence increased the cumulative odds of extinction 1.5-fold and 2.0-fold for high-and low-diversity populations, respectively (Fig. 3*Ciii*,*iv*, Table F1). Greater initial maladaptation (i.e., a mean initial genotype further from θ) was associated with increased extinction risk across all treatments (Fig. 3*D,* Table F1). The effect of maladaptation on extinction was stronger for large populations than small populations (Table F1); in large populations with high diversity, only the most strongly maladapted populations had considerable extinction risk (Fig. 3*Di*). Density dependence increased extinction risk for all genotypes and only marginally affected the relationship between genotype and maladaptation (i.e., raised the intercept while slightly lowering the slope, Fig. 3*D,* Table F1). The combined result was that in large populations with density-independent growth, only populations that were highly maladapted were at risk of going extinct, but under density-dependent growth all populations had substantial extinction risk (Fig. 3*D*).

**Figure 3:**
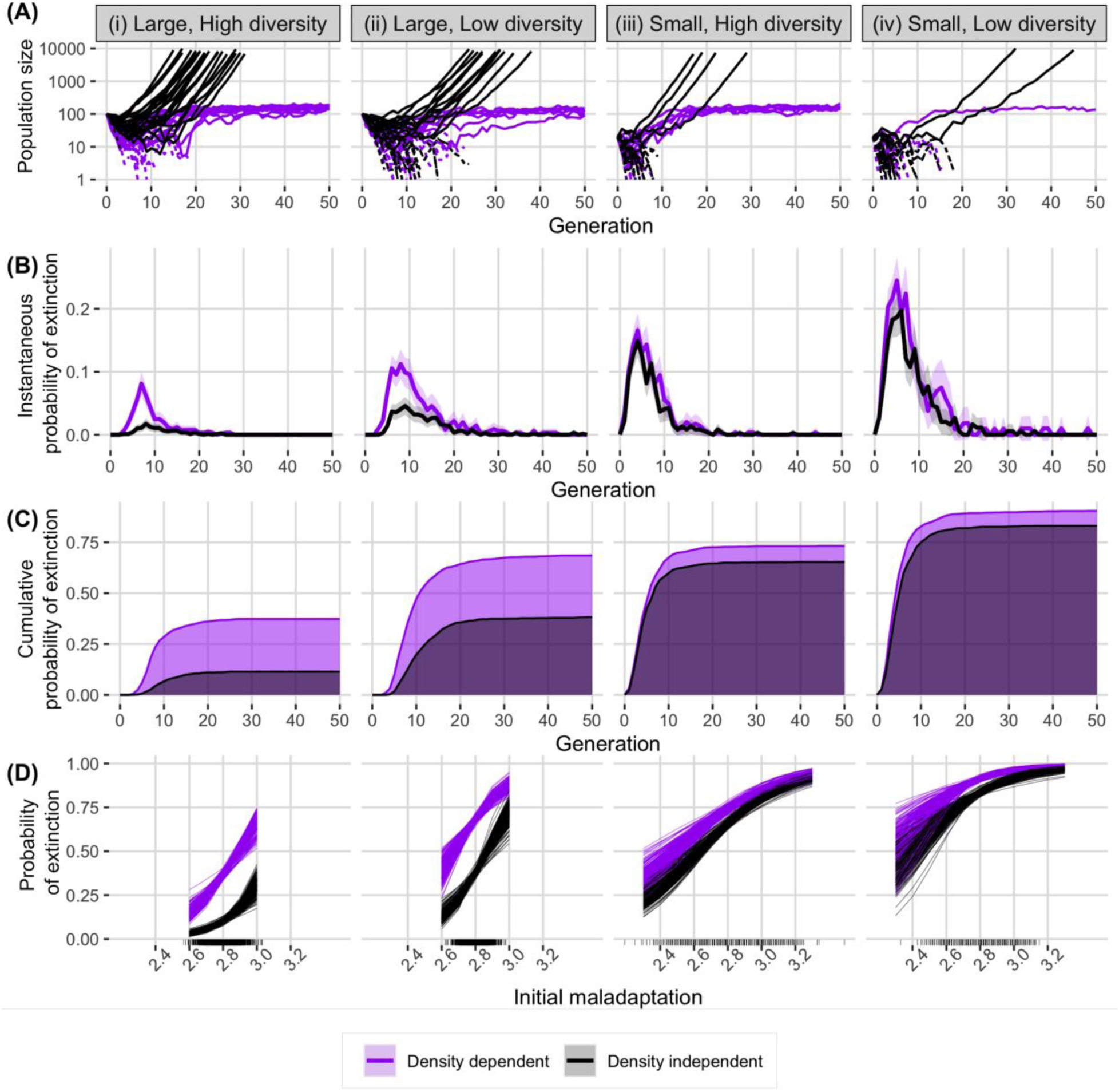
Population size and extinction characteristics of 1000 simulations per treatment group run for up to 50 generations. (A) mean population size of 25 randomly selected trials for each treatment. Dashed lines are extinct populations. Simulations were truncated for populations surpassing size 10000. (B) Instantaneous probability of extinction in each generation, conditioned on survival until that generation. Ribbons represent twice the standard error on both sides of the estimate. (C) Cumulative probability of extinction over time. Areas under the curve are shaded to highlight the difference between density-dependent and density-independent populations. (D) 200 posterior estimates per treatment of extinction probability as a function of initial degree of maladaptation, as predicted by a generalized linear model (Table 2). Marks on x-axis indicate the distribution of initial mean population genotypes for those treatments.

The results for probability of rescue were qualitatively identical to those for probability of extinction but with directions switched (Table 1, Table F2, Figure F1). Likewise, the results were qualitatively similar regardless of the type of rescue (i.e., fitness-based rescue or size-based rescue). For both types, the probability of rescue declined with density dependence and density dependence had stronger effects in large populations than small ones. Density dependence increased mean rescue time only slightly (less than 1.5 generations) for all population treatments except for large populations, for which density dependence increased the time to return to initial size by 9-10 generations (Fig. F2). Notably, many small populations were rescued instantaneously, i.e., they achieved three generations either exceeding initial size or 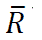 > 1 (or both) either instantly or after one time step (Table 1, Fig. 3*A*). These “instantaneous rescues” happened primarily in small populations, comprising up to 16% of successful rescues observed in small population treatments. Both the overall frequency and the proportion of total rescues comprising these instantaneous rescues decreased with low genetic diversity and with density dependence. Pooling across all large initial size treatment populations, only four populations achieved instantaneous rescue of either type, comprising less than 1% of rescue episodes for all large treatments. Extinctions occurred after both fitness-and size-based rescue (Table 1). Extinction after rescue was more common for small populations than large ones, for low-diversity populations than high-diversity ones, and under density dependence than density independence for both size-and fitness-based rescue.

**Table 1.** Summary of rescue probabilities, proportion of rescue events occurring instantly, times to rescue, and extinction probabilities after rescue 50 generation simulations for both types of rescue.

| Size and diversity | Prob. rescue |  | Prop. (number) instant rescue† |  | Mean time to rescue |  | Prob. extinct after rescue |  |
| --- | --- | --- | --- | --- | --- | --- | --- | --- |
|  | DI | NDD | DI | NDD | DI | NDD | DI | NDD |
| Fitness-based rescue |  |  |  |  |  |  |  |  |
| Large, high | 0.92 | 0.71 | 0.00 (3) | 0.00 (0) | 5.1 | 6.7 | 0.04 | 0.12 |
| Large, low | 0.69 | 0.41 | 0.00 (0) | 0.00 (0) | 8.3 | 9.2 | 0.10 | 0.24 |
| Small, high | 0.44 | 0.36 | 0.16 (68) | 0.06 (23) | 4.0 | 4.7 | 0.21 | 0.27 |
| Small, low | 0.24 | 0.16 | 0.04 (11) | 0.01(2) | 5.7 | 7.0 | 0.31 | 0.39 |
| Size-based rescue |  |  |  |  |  |  |  |  |
| Large, high | 0.89 | 0.62 | 0.00 (1) | 0.00 (0) | 11.2 | 20.3 | 0.00 | 0.00 |
| Large, low | 0.62 | 0.26 | 0.00 (0) | 0.00 (0) | 17.4 | 27.7 | 0.00 | 0.00 |
| Small, high | 0.37 | 0.29 | 0.16 (61) | 0.05 (15) | 6.3 | 7.9 | 0.06 | 0.07 |
| Small, low | 0.19 | 0.12 | 0.13 (24) | 0.08 (9) | 7.9 | 8.6 | 0.12 | 0.17 |

† proportion instant rescue is the proportion of rescued populations in the treatment where rescue occurred in 0 or 1 generations; the number is the total number of observed populations (out of 1000) in the treatment experiencing instantaneous rescue

### Genetic variance, fixation, and fitness

Over the first 15 generations, mean additive genetic variance, 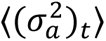, declined for all treatments (Fig. 4). Surviving density-dependent populations lost more genetic diversity than density-independent populations (Fig. 4, solid black *vs.* solid purple lines); this effect was greater in large populations. Across treatment and throughout time, populations that went extinct had lower mean genetic variance than populations that survived (Fig. 4, solid *vs.* dashed lines). This loss of genetic variation was associated with fixation for both positive (adaptive) and negative (maladaptive) alleles (Fig. G1). Populations that went extinct typically had on average more loci at fixation for either allele than surviving populations, although in large populations this difference did not appear until the fifth generation, when populations approached minimum size (Fig. G1). Differences in the trajectories over time of genotype and intrinsic fitness between density-dependent and density-independent populations conditioned on survival or extinction were slight (Figs. G2,G3).

**Figure 4:**
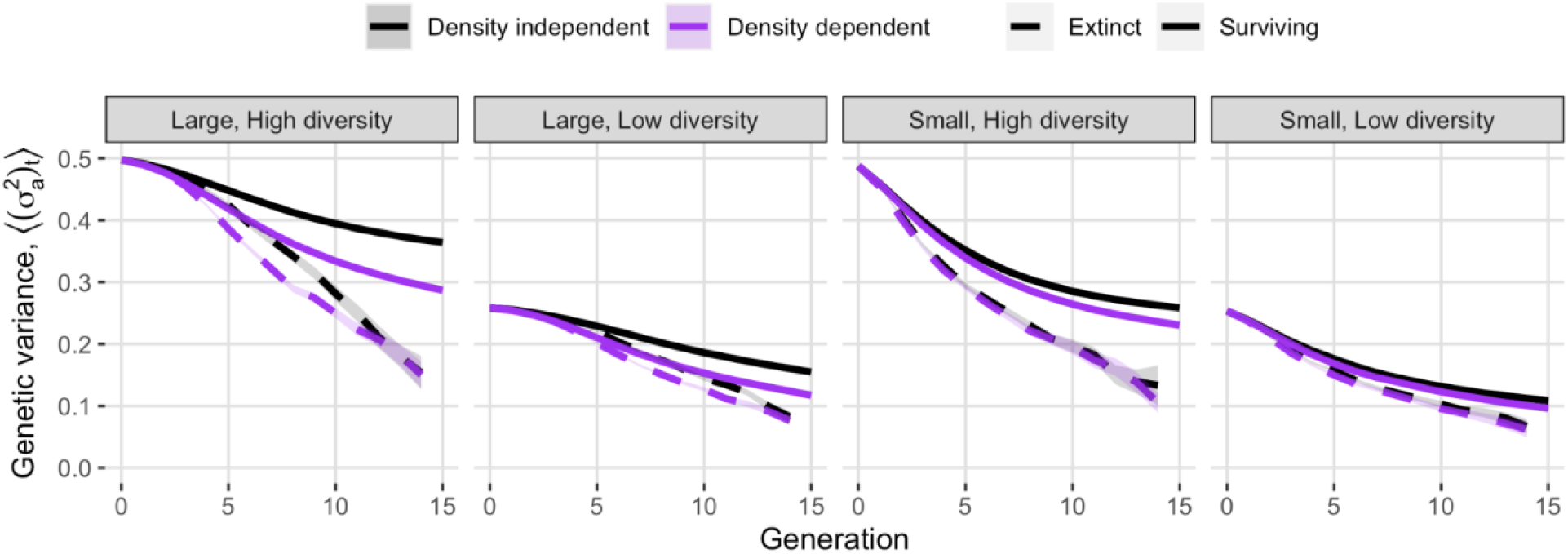
Mean additive genetic variance in surviving and extinct populations. Shaded areas represent twice the standard error on each side of the mean (the standard error is often too small to be visible on this plot).

### The extinction vortex

Extinct populations in all treatments within the 15-generation simulations had negative mean population growth rate (⟨*r*_τ_⟩, Fig. 5*A*) and non-zero mean loss of genetic diversity (⟨ν_τ_⟩, Fig. 5*B*) in each time step. Declines in population growth rates and rate of loss of genetic diversity accelerated, as expected for an extinction vortex [25,34] in the three generations prior to extinction (*τ* ≤ 3), with populations losing an average of 17% of their standing additive genetic variance in the final generation preceding extinction. In contrast, the rate of adaptation (1 − ⟨κ_τ_⟩) decreased steadily over time in all treatments rather than accelerating immediately prior to extinction (Fig. 5*C*). In the three generations prior to extinction trends in density-dependent and density-independent populations were similar. Four or more generations prior to extinction (*τ* > 3), large density-dependent populations had faster population decline (i.e., more negative ⟨*r*_τ_⟩) than large density-independent populations, for which the rate of population decline was constant over time (Fig. 5*A*). In contrast, density-dependent and density-independent populations had similar rates of loss of genetic diversity (all differences in ⟨ν_τ_⟩ between density-dependent and density-independent treatments within 0.02, Fig. 5*B*). In the generation prior to extinction, populations in all treatments had a mean adaptation rate of approximately zero (i.e., ⟨κ_τ=1_⟩ ≈ 1 and ⟨κ̅_τ=1_⟩ ≈ ⟨κ̅_τ=2_⟩). Figures of non-normalized state variables conditioned on generation of extinction, revealing qualitatively similar results, are available in Supporting Information H.

**Figure 5:**
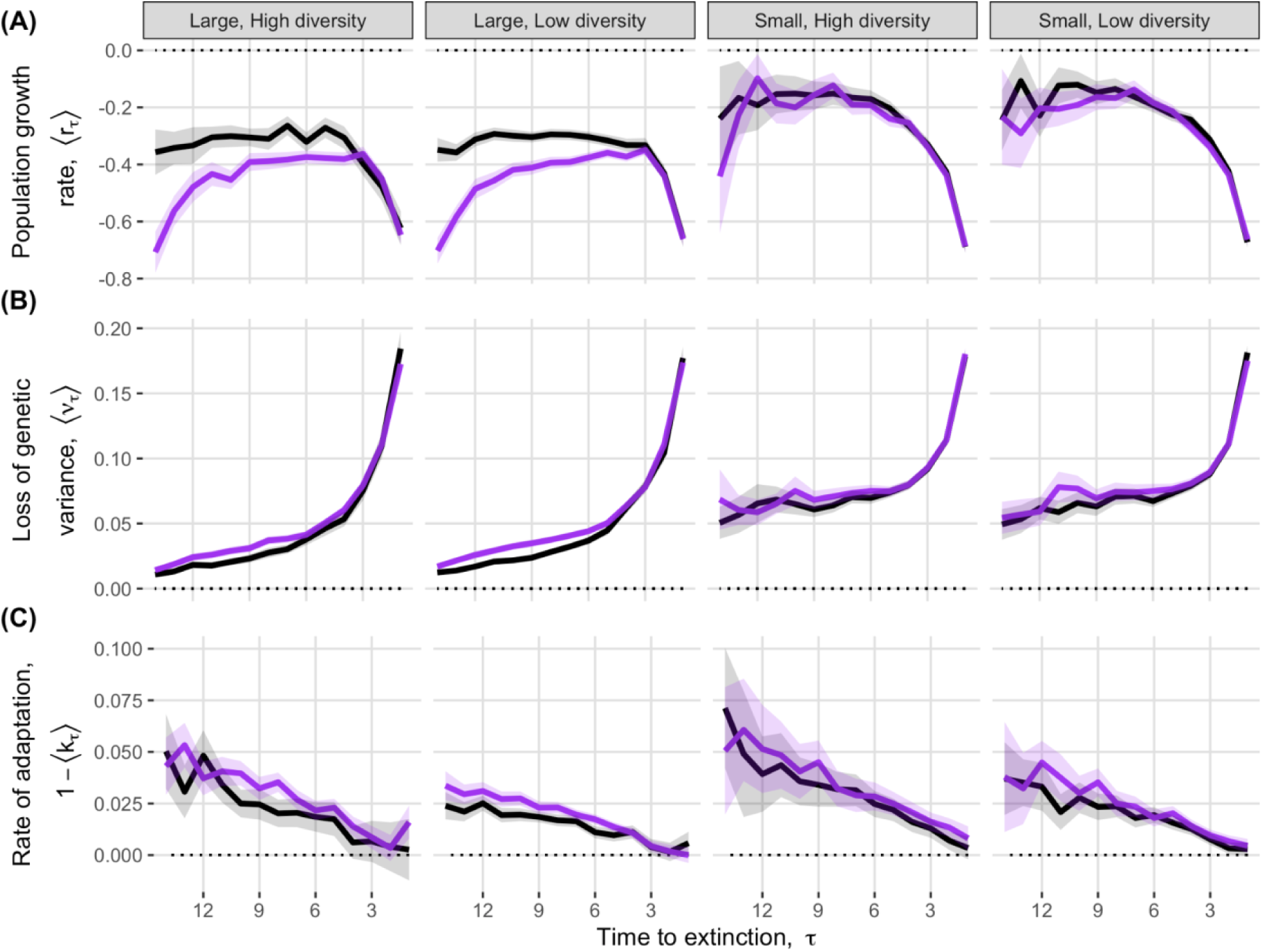
Rates of change as a function of time to extinction (τ), demonstrating the presence of an extinction vortex, in (A) population growth, (B) loss of genetic variation, and (C) rate of adaptation. Small τ represents time steps closer to the time of a population’s extinction. Shaded areas represent twice the standard error on both sides of the mean.

## Discussion

We found that evolutionary rescue is constrained by the combined effects of negative density dependence and loss of genetic variation. For all treatments, negative density dependence decreased population growth rates, causing populations to decline more quickly, reach lower densities, and stay at low densities for longer. These effects fell disproportionately on large populations, which previous theory predicted to be at low extinction risk under changing environments. As populations declined, they lost additive genetic variance and faced slowing rates of adaptation. However, declining genetic variance had contrasting effects on density-independent versus density-dependent populations, increasing growth rates under density-independent growth and decreasing them under density dependence, respectively. These demographic and genetic processes often reinforced each other to produce an extinction vortex, precluding evolutionary rescue. Here, we discuss the effects of negative density dependence, loss of genetic variation, and the extinction vortex on evolutionary rescue.

### Interactive effects of density dependence and genetic erosion

We found an important interaction between size and density dependence not noted in previous models of rescue. Theory [1] and empirical studies [3–5] predict that in density independent settings, initial population size confers an advantage to populations experiencing environmental change; our model agrees with this finding (Fig. 3, Table F1). However, we found that density dependent growth considerably reduced the advantage that initial size conferred (Fig. 3, Table F1). If the density dependence produces a carrying capacity, upon environmental change populations at or near this carrying capacity will experience more rapid decline and reach lower densities than density-independent models predict. Natural populations and conservation targets may be below carrying capacity due pressures such as predation [35] or inbreeding depression [36]. While populations considerably below carrying capacity are at a disadvantage due to increased demographic stochasticity, their growth rates will be less affected by density dependence. Thus, populations considered at low risk by classic, density-independent models may face a non-trivial threat of extinction from environmental change under more realistic density-dependent population growth.

Adaptation to environmental change was accompanied by a loss of genetic diversity and potential for adaptation not captured in most models (but see [23,24]). Quantitative genetic models often assume populations are large for analytical tractability [10]. Under this assumption, fixation and loss of alleles are rare as there is only marginal sampling variation and thus minimal drift. However, population decline is by definition part of the rescue process. Our model demonstrates that additive genetic variance is lost even in populations that survive the environmental change (Fig. 4). This loss of genetic diversity was associated with slowing rates of adaptation, contributing to prolonged time until rescue. This loss was partly driven by fixation of both adaptive and maladaptive alleles, typically occurring when populations were at their smallest size (Fig. G1). This matches theoretical results [23] as well as empirical results in a seed beetle adapting to a novel host plant [37]. Likewise, laboratory studies with *Tribolium castaneum* adapting to novel conditions found that accumulating genetic load likely slows the rate of adaptation [38] and that more genetic diversity likely assists population recovery through higher rates of recombination [4]. We therefore recommend that evolutionary rescue models include genetic erosion where appropriate, as environmental stress [39] and inbreeding due to bottleneck events [11] are known to erode standing genetic variation [12].

Our simulation results also showed a surprising interaction between density dependence and genetic erosion. Under density-dependence, the effects of genetic erosion were negative, although small, and did not emerge until nearly the end of simulations (Fig. 2). Conversely, under density independence, populations that experienced genetic erosion had higher growth rates than populations with genetic diversity held constant. In total, this meant that the effect of density dependence on final population size was considerably larger under genetic erosion than in the analytical model, where genetic diversity was held constant. While theory primarily focuses on the effects of genetic and phenotypic diversity on the rate of adaptation, there is a second effect evident in equation (2) that is often ignored in analysis of rescue: a “variance load” [16,40] where mean population fitness is decreased by the increasing the mean phenotypic distance from the optimum. Our results can be explained by noting that adaptation in response to selection reduces phenotypic variance and thus reduces variance load. In populations reaching low densities due to density dependence, the positive effect of reduced variance load on population growth is counteracted by drift (including the fixation of maladaptive alleles, Fig. G1*B*) and increased extinctions (Fig. 3), such that genetic erosion has a net negative effect on expected population size. However, under density independence, populations are less likely to reach low densities and experience drift, meaning the net effect of reduced phenotypic variance is positive, as it is associated with adaptation. This further highlights the need to simultaneously include negative density dependence and effects of stochasticity at small densities in models of rescue.

### How an extinction vortex impedes evolutionary rescue

Challenging environments can draw populations into an extinction vortex [25]. Although previous analyses have hypothesized that reduced genetic diversity associated with declining population size may reduce adaptive potential [18,21], to our knowledge our analysis is the first to demonstrate the existence of positive feedback loops connecting declining genetic diversity, reduced rates of adaptation, and reduced population growth that characterize this vortex ([25], Figs. 1,5 in the present study) in the context of evolutionary rescue. Gilpin and Soule’s original conception of the extinction vortex [25] did not assume a population is severely maladapted to its environment, only that small density increases susceptibility to drift, inbreeding, and demographic stochasticity. In contrast, evolutionary rescue is defined by populations facing severe maladaptation that pushes mean fitness below the rate of replacement. Thus, rather than an extinction vortex arising by a population slowly becoming less fit due to drift, a population experiencing environmental change immediately becomes less fit and also faces an extinction vortex *via* drift and stochasticity that further reduces its ability to sufficiently adapt to the new environment (Fig. 5*B,C*). Negative density dependence, rather than changing the demographic and genetic dynamics in the generations immediately preceding extinction, “widens” the opening to the vortex as populations are more likely to quickly decline to small size several generations before extinction (Fig. 5*A*). For managers of small, at-risk populations that are susceptible to sudden environmental change and extinction, alternative strategies such as assisted migration may be needed to pull populations out of the self-reinforcing demographic and genetic processes of the vortex [5,41].

### Rapid rescue in small populations

Upon environmental change, some small populations already had a large proportion of phenotypes adapted to the environmental change due to sampling variation, allowing them to be rescued very quickly (Fig. 3*D*, Table 1). In contrast, large populations were rarely rescued immediately (Table 1) because the average initial phenotype was less variable across populations (Fig. 3*D*). These “rapid rescues” appear to be a polygenic analog of the phenomenon demonstrated in a haploid single-locus model [22], where small populations may, under certain conditions, have an overrepresentation of individuals with genotypes adapted to the novel environment. In our quantitative trait model where adaptive and maladaptive alleles have equal effect on the genotype, small populations are just as likely to have an over-representation of especially maladapted genotypes as they are to have an over-representation of favored genotypes (Fig. 3*D*). Another interpretation of this result is the fact that a sizable proportion of small populations were rescued instantly due to a “lucky” initial genetic state (Table 1) underscores the difficulties that small populations without this “luck” have in adapting to novel environments.

### Operational use of current rescue definitions

Current definitions of rescue are difficult to operationalize in stochastic settings. Rescued is often understood to mean “not extinct”, but this definition is lacking, as our analysis shows that even under relatively short timescales (50 generations in our simulations), populations can show sensible indications of “rescue” and still go extinct (Table 1). Likewise, in conservation or management settings, quantifying fitness to determine when mean fitness (intrinsic or realized) exceeds unity may not be feasible [6]. Genetic or demographic stochasticity (to say nothing of environmental stochasticity or other sources of variation) complicate this further, as some natural benchmarks such as population increase may easily be undone causing populations to revert to an “unrescued” state. We defined rescue with two heuristic criteria which provided qualitatively similar (although not identical) results (Table 1), but they were not guaranteed to be simultaneously met in the same population. Having more readily operationalizable definitions of rescue could prove to be important both for empiricists testing theoretical predictions and for managers and conservation practitioners seeking to use the theory.

### Implications for conservation and management

Our simulations were on short timescales relevant to immediate or near-term conservation concern. However, we note that the reductions in population-size or genetic diversity may be important on longer timescales, even after populations have been rescued. An effective population size on the order of hundreds is needed to reduce longer-term extinction risk due to accumulation of deleterious alleles [42–44]. Populations need to spend long durations at large size to offset even brief episodes of drift and genetic erosion [43], like those that might be seen during the “trough” or low point in population size following environmental change. Under density dependence, not only are recovered populations held at carrying capacity, potentially precluding this re-establishment of variation, but they also spend more time at smaller densities before being rescued (Figs. 2,3*A*), leading to greater genetic erosion (Fig. 4). Indeed, our simulations showed that up to 40% of populations in a treatment group that were “rescued” under our criteria go extinct within 50 generations, with more extinctions in populations with initially low genetic diversity (Table 1). Furthermore, population size and remaining standing genetic variation are important for persisting through future environmental changes. The erosion of genetic variation is similar to a genetic bottleneck that would hinder a population’s ability to adapt to additional environmental changes [11,45]. In our simulations, we observed fixation of the positive allele in surviving populations (Fig. G1*A*); in the event of future environmental change favoring another allele, its loss following the first environmental change event would hinder subsequent adaptation [2,20].

Here, we introduce a biologically realistic model for a more complete and applicable understanding of population rescue after environmental change, providing better prediction of potential outcomes and guidance for conservation and management as well as providing testable predictions for empirical validation. We demonstrate that negative density dependence impedes persistence in a novel environment, and that large and well-adapted populations previously predicted to be at minimal extinction risk [1,3,24] do, in fact, face non-trivial risk of extinction under density dependence. For managers, one prescription is to increase habitat carrying capacities, for example by increasing resource availability, which will alleviate reductions in growth rates as populations recover from environmental change.

Although our simulated experimental design only featured density dependence as a binary treatment, in natural settings it is a continuous variable that managers may be able to influence. Because expected time to extinction increases exponentially with carrying capacity [17], decreasing strength or degree of competition could drastically improve long-term population fates, particularly for large populations previously perceived as having low extinction risk. A corollary is that, in addition to other predictors of extinction risk under rescue [3,7], the carrying capacity or degree of intraspecific competition should be considered when assessing risks to populations subject to novel habitat alteration, degradation, or reduction.

Furthermore, managers implementing strategies to maximize evolutionary potential [45] should consider how competition and population size may influence adaptation under environmental or climate change. These predictions highlight the effects of density dependence and loss of genetic diversity on populations exposed to severe environmental change, which is crucial in an age of global anthropogenic changes that put populations at unprecedented risk of extinction.

## Acknowledgements

The authors would like to thank Robert Holt for helpful suggestions following an early presentation on work from this project and Ophélie Ronce for comments that improved the manuscript. This work was supported by NSF award 1930222 to BAM and 1930650 to RAH. SWN was also supported by NSF IGERT award 114807 while working on this project and LFD was also supported by NSF Graduate Research Fellowship number 006784.

## Data and code availability

Data and simulation code to produce data and analysis will be archived on Dryad upon acceptance of the manuscript. Code and output files are also stored on GitHub at the address https://github.com/melbourne-lab/evo_rescue_ndd_erosion.

