## Supporting Information, A-H for "How density dependence, genetic erosion, and the extinction vortex impact evolutionary rescue"

**Supporting Information A: Derivation of additive genetic variance**

The breeding value of individual *i* at locus *j*, $b_{ij}$, is defined as twice *i*'s deviation from the population mean genetic value at locus *j* (Lynch and Walsh 1988, pp. 72 – 73). Note that the breeding value is a mean-centered genetic value, i.e., $b_{ij} = g_{ij} - \mu_j$ where

$$g_{ij} = \begin{cases} \frac{1}{\sqrt{m}}\ if\ i\ is\ homozygous\ for\ positive\ allele\ at\ locus\ j \\ 0\ if\ i\ is\ heterozygous\ at\ locus\ j \\ \frac{-1}{\sqrt{m}}\ if\ i\ is\ homozygous\ for\ negative\ allele\ at\ locus\ j \end{cases}$$

and $\mu_j$ is the mean genotypic value across all individuals in the population at locus *j.* According to Lynch and Walsh (1998, p. 78), when breeding values are purely additive (i.e., there is no dominance or epistasis) and when mating is random, variance in breeding values at one locus within a population is equivalent to the additive genetic variance in the population at that locus. Because the genetic value is the breeding value plus the population-level mean genotype at that locus (a constant), genetic values and breeding values have the same variance, and thus variance in genetic values at locus *j* is equivalent to the additive genetic variance at locus *j*.

Define $p_j$ as the frequency of the positive allele at locus *j.* Individual *i*'s genetic value at locus *j* is a Binomial random variable scaled by $\frac{1}{\sqrt{m}}$ and shifted by $\frac{-1}{\sqrt{m}}$, with n (the number of trials) equal to 2 and probability $p_j$. Thus, the population variance in $g_{ij}$ (and therefore the additive genetic variance at locus j, $\sigma_j^2$) is equal to

$$\left(\frac{1}{\sqrt{m}}\right)^2 2p_j(1-p_j) = \frac{2p_j(1-p_j)}{m}.$$

Loci are segregated independently and therefore distributed independently. An individual's genotype is the unweighted mean of genetic values across loci; therefore, the variance in genotypes is the sum of variances at individual loci, i.e.,

$$\sigma_a^2 = \sum_{j=1}^{m} \frac{2p_j(1-p_j)}{m} = 2\left(\frac{1}{m}\sum_{j=1}^{m} p_j - \frac{1}{m}\sum_{j=1}^{m} p_j^2\right) = 2\left(p' - \frac{1}{m}\sum_{j=1}^{m} p_j^2\right)$$

where p' is the arithmetic mean of allele frequencies across all loci. The remaining summation is the expectation of squared allele frequencies. Because $V[p] = E[p^2] - E[p]^2$, the summation is equivalent to $\sigma_p^2 + p'^2$ where $\sigma_p^2$ is the variance across loci in allele frequencies. Thus, the above expression can be rewritten as

$$\sigma_a^2 = 2(p' - p'^2 - \sigma_p^2) = 2p'(1 - p') - 2\sigma_p^2.$$

The first term in the final expression is variance associated with the diversity of alleles and the latter term is sampling variance and is analogous to drift.

In this model, an individual's phenotype is a normally distributed random variable with mean equal to its genotype and variance equal to a constant $\sigma_e^2$. This means that total phenotypic variance is $\sigma_a^2 + \sigma_e^2$ and heritability is $h^2 = \sigma_a^2 / (\sigma_a^2 + \sigma_e^2)$ (Lynch and Walsh 1998).

**Supporting Information B**

**Table B1** Parameter and variable definitions for the model.

| Symbol | Definition | Value used or range |
|---|---|---|
| | Fixed parameters | |
| $m$ | Number of bi-allelic loci | 25 |
| $a$ | Strength of density dependence | 0 (density independent) |
| | | 0.0035 (density dependent) |
| $N_0$ | Initial population size | 20 (small population) |
| | | 100 (large population) |
| $w^2$ | Width of fitness landscape | 3.5 |
| $\sigma_e^2$ | Environmental component of phenotypic variance | 0.5 |
| $\theta$ | Optimal phenotype | 2.8 |
| | Individual-level variables | |
| $g_i$ | Genotype of individual *i* | $[-1/(2\sqrt{m}), 1/(2\sqrt{m})]$ |
| $z_i$ | Phenotype of individual *i* | $[-\infty, \infty]$ |
| $W_i$ | Intrinsic fitness of individual *i* | $[0, W_{max}]$ |
| $R_i$ | Expected number of offspring for individual *i* | $[0, W_{max}]$ |
| | Population-level variables | |
| $p_j$ | Frequency of the positive allele at locus $j$ | (0, 1) |
| $p'$ | Mean frequency of positive alleles across all loci | (0, 1) |
| $N_t$ | Population size at time $t$ | $[0, \infty)$ |
| $\sigma_{at}^2$ | Additive genetic variance among individuals at time *t* | [0, 0.5] |
| | Derived population-level variables | |
| $r_\tau$ | (Log) mean population growth rate $\tau$ generations before extinction | $(-\infty,\infty)$ |
| $\nu_\tau$ | Loss of previous generation's additive genetic variation $\tau$ generations before extinction | [0, 1] |
| $1 - k_\tau$ | Rate of genotypic adaptation $\tau$ generations before extinction | [0, 1] |

**Supporting Information C: Justification of parameter values and notes on model parameterization**

By Equation (1) in main text, the maximum additive genetic fitness a population can attain is $\sigma_a^2 = 0.5$, for the special case where all loci have $p_j$ = 1/2 across the population (in which case $\sigma_p^2 = 0$). Our populations had initial additive genetic variances of $\sigma_a^2 \approx 0.5$ and $\sigma_a^2 \approx 0.25$ for high- and low- diversity populations, respectively. We selected a non-genetic phenotypic variance parameter, $\sigma_e^2$, value of 0.5 to produce initial heritabilities resembling fitness heritability estimates obtained for *Tribolium castaneum* by Wade et al. (1996) in a controlled laboratory setting. Likewise, we set a maximum intrinsic fitness value of $W_{max} = 2$ based on prior work with *T. castaneum* finding maximum population growth rates of approximately 2 in ideal, density-independent laboratory conditions (e.g., Hufbauer et al. 2015). We used a fitness landscape with width $w^2$ = 3.5 ($w \approx 1.87$); we justify this value below.

We note here that the dynamics of phenotypic change over time in this model can be described in part by the ratio $\gamma^2 = \sigma^2/w^2$ , where $\sigma^2 = \sigma_a^2 + \sigma_e^2$ is the overall phenotypic variance in the population. The case of $\gamma^2$ approaching infinity corresponds to a case where there is high phenotypic variance in the population and selection pressure is very strong. The case of $\gamma^2$ approaching zero corresponds to relatively low phenotypic variance and very weak selection. We demonstrate below that the conditions under which evolutionary rescue may or may not occur (i.e., cases where extinction or persistence under novel environmental change are not trivially likely outcomes) are confined to a narrow range of $\gamma^2$ values.

In the model of Gomulkiewicz and Holt (1995), the log population size in generation $t$ is described by the following equation:

$$n_t = n_0 + t[\hat{w} + (1/2)\, log\, (w^2/(w^2+\sigma^2))] - (1/2)\, \underline{z}_0^2/(w^2+\sigma^2)\, [(1-k^{2t}) / (1-k^2)],$$

where $n_0$ is the log of the initial population size, $\hat{w} = log\,(W_{max})$ , $\underline{z}_0$ is the initial population mean phenotype, and $k = (w^2 + (1-h^2)\sigma^2) / (w^2+\sigma^2))$ is the amount of phenotypic variance remaining after one round of selection given heritability $h^2$. Substituting in $\gamma^2$ in for $w^2$ and $\sigma^2$ gives

$$n_t = n_0 + t[\hat{w} - (1/2)\, log\,(1+\gamma^2)] - (1/2)\,(\underline{z}_0/w)^2 / (1+\gamma^2)\, [(1-k^{2t}) / (1-k^2)],$$

and $k$ can be written as $k = (1 + (1-h^2)\gamma^2) / (1+\gamma^2))$. We note that this parameterization is analogous to a model where phenotypes are scaled by the width of the selection landscape; in this case, scaled phenotypes are normally distributed with mean $\underline{z}_0/w$ and the phenotypic variance is $\gamma^2$.

The second term in this sum is the maximum growth rate minus a demographic penalty for standing genetic variation ("variance load") in the population. Because the third term in the sum is negative, this second term must be positive for population growth to occur in the long-term. For a population with a maximum intrinsic fitness on the order of 1 (e.g., in our case maximum intrinsic fitness is 2), $\hat{w}$ will be on the order of 0. As $\gamma^2$ grows larger than order 1, the variance load term overwhelms the maximum growth rate, $\widehat{w}$, due to an increasing phenotypic variation subject to harsher selection, contributing to population decline increasing linearly in time (i.e., producing exponential decay). This demonstrates that for persistence and evolutionary rescue to occur, $\gamma^2$ can not be on a larger order of magnitude than $W_{max}$ . In our simulations, we set $W_{max} = 2$, necessitating that $\gamma^2$ can not be much larger than 1 (i.e., $\sigma^2$ and $w^2$ must be on similar orders of magnitude).

The third term in the sum above is "evolution load" (Lande and Shannon 1996) caused by phenotypic mismatch to the environment. Assuming $\gamma^2$ < 1 (due to the upper bound specified above), here we demonstrate that $\gamma^2$ also must have a lower bound for non-trivial dynamics to occur. The contributions of evolution load to decline shrink over time by a factor of $k^2$ (note here that the term $(1-k^{2t})/(1-k^2)$ is a partial sum of $k^2$), where $k^2$ close to 1 produces very gradual phenotypic change. The proportion $k^2$ can be expressed as

$$k^2 = [1 + 2\gamma^2(1-h^2) + (\gamma^2(1-h^2))^2] \,/\, [1 + 2\gamma^2 + (\gamma^2)^2].$$

As $\gamma^2$ approaches zero, this term approaches 1, i.e., no adaptation (because there is weak selection and relatively little phenotypic variation), suggesting that for adaptation to happen on a rapid timescale (e.g., 15 generations as we model), $\gamma^2$ must have a lower bound. For $\gamma^2$ smaller than order (0.1), $(\gamma^2)^2$ becomes negligible, suggesting that $\gamma^2$ on the order of 0.1 is a reasonable lower bound. If $\gamma^2$ is too small (e.g., $\gamma^2 < 0.1$), then adaptation will proceed very slowly. In this case, the population dynamics will be dominated by the initial condition. This can be illustrated by examining the limiting case for $n_t$ of $\gamma^2$ set to zero:

$$n_t = n_0 + (\widehat{w} - (1/2)(\underline{z}_0/w)^2)t.$$

Here, because there is no phenotypic variance or adaptation, the population dynamics are determined entirely by the maximum growth rate and the maladaptation load. Considering that evolutionary rescue is predicated on populations facing an environmental change that pushes mean population fitness below replacement levels, the most likely outcome is extinction.

From this we can conclude that evolutionary rescue is most likely to occur for $\gamma^2$ between 0.1 and 1, and considerably less likely to occur outside of this range due to being dominated by variance load for larger $\gamma^2$ and the population not adapting (and going extinct or persisting entirely due to the magnitude of its original phenotypic load) for larger $\gamma^2$. This means that evolutionary rescue is restricted primarily to cases where the phenotypic variance on short-to-medium timescales is 0.1-1 times the width of the fitness landscape. Our initial phenotypic variances (approximately 0.75 for the low diversity populations and 1 for the high diversity populations) are 0.21 and 0.28 times the width of the fitness landscape that we use ($w^2$ = 3.5).

**Supporting Information D: Relationship between phenotypic optimum and allele frequencies**

If the mean frequency of the positive allele across all loci is $p'$, then the mean genotype is

$$\underline{g} = \frac{2mp'}{2\sqrt{m}} - \frac{2m(1-p')}{2\sqrt{m}} = \sqrt{m}\,(2p' - 1).$$

Consequently, genotypes are bounded between $-\sqrt{m}$ and $\sqrt{m}$. Taking the inverse of the above expression gives the mean positive allele frequency as a function of the optimum genotype,

$$p' = \frac{1}{2} + \frac{\underline{g}}{2\sqrt{m}}.$$

Multiplying both sides of this equation by the total number of allele copies, $2m$, gives the mean number of positive alleles per individual in the population. In our implementation of this model with *m = 25* loci; populations with an initial mean genotype of *p'* = 0.5 and therefore individuals have on average 25 copies of the positive allele. By the above relationship, an environmental shift with phenotypic optimum (and therefore genotypic optimum, $\underline{g}$) of θ = 2.8 favors a genome with 39 copies of the positive allele per individual. Thus, 14 alleles must change from negative to positive for the typical population to reach the new phenotypic optimum.

**Supporting Information E: Description of Bayesian models and estimation of effect sizes**

We fit separate models for extinction and two types of rescue (see Methods) using Bayesian GLMs fit in the R package rstanarm, version 2.21.1 (Goodrich et al. 2020). For all three models, we used 1000 simulations, each of up to 50 generations, per parameter combination (n = 8000 total trials). Treatments (density dependence, initial size, initial genetic diversity) were modeled as categorical variables and maladaptation was modeled as a continuous variable. In these models, the intercept for each treatment group is the mean log-odds of the response (extinction or rescue) for a population with mean initial maladaptation equal to θ (i.e., mean population genotype of zero). Each parameter was defined by a normally distributed prior with mean of 0 and standard deviation of 5. All models were run for four chains, each of 2000 iterations with a warmup period of 1000 iterations, giving 4000 posterior estimates per parameter. All models converged with Rhat for each parameter approximately 1.0.

We estimated the effect of changing a single parameter on the response while holding all other parameters constant by estimating the differences in intercepts. We estimated an intercept for each parameter combination for each posterior draw; thus, credible intervals for each effect size account for covariances among model parameters. Models were fit with baseline case (intercept) of large, high diversity, density-independent populations. As such, effect sizes for each parameter give the respective influences of decreasing population size, decreasing genetic diversity, or adding density dependence, respectively.

**Supporting Information F: Analysis of probability of extinction and rescue**

**Table F1:** Effect sizes of treatment variables changed while others are held constant on log odds of extinction in 50 generations and accompanying 95% credible intervals.

| Variable changed | Held constant | Effect size (95% CI) |
|---|---|---|
| Density independent to dependent | Large, high diversity | 1.60 (1.36, 1.85) |
| | Large, low diversity | 1.27 (1.08, 1.46) |
| | Small, high diversity | 0.41 (0.21, 0.61) |
| | Small, low diversity | 0.71 (0.44, 1.00) |
| Large to small | High diversity, DI | 2.82 (2.58, 3.08) |
| | High diversity, NDD | 1.63 (1.44, 1.83) |
| | Low diversity, DI | 2.13 (1.92, 2.35) |
| | Low diversity, NDD | 1.58 (1.31, 1.85) |
| High to low diversity | Large, DI | 1.67 (1.43, 1.91) |
| | Large, NDD | 1.34 (1.15, 1.53) |
| | Small, DI | 0.99 (0.77, 1.21) |
| | Small, NDD | 1.29 (1.03, 1.56) |
| Increased maladaptation | Large, high variance, DI | 6.65 (4.38, 8.99) |
| | Large, high variance, NDD | 6.04 (4.16, 7.99) |
| | Large, low variance, DI | 7.24 (4.77, 9.79) |
| | Large, low variance, NDD | 6.19 (3.82, 8.55) |
| | Small, high variance, DI | 3.76 (2.90, 4.63) |
| | Small, high variance, NDD | 3.39 (2.40, 4.40) |
| | Small, low variance, DI | 3.99 (2.46, 5.50) |
| | Small, low variance, NDD | 4.48 (2.67, 6.34) |

**Table F2:** Effect sizes associated with the probability of fitness- and size-based rescue within 50 generations, according to Bayesian GLMs.

| Variable changed | Held constant | Fitness rescue effect size (95% CI) | Size rescue effect size (95% CI) |
|---|---|---|---|
| Density independent to dependent | Large, high diversity | -1.55 (-1.83, -1.28) | -1.66 (-1.90, -1.42) |
| | Large, low diversity | -1.16 (-1.34, -0.97) | -1.55 (-1.74, -1.36) |
| | Small, high diversity | -0.35 (-0.53, -0.17) | -0.41 (-0.60, -0.22) |
| | Small, low diversity | -0.58 (-0.82, -0.34) | -0.65 (-0.92, -0.38) |
| Large to small | High diversity, DI | -2.75 (-3.02, -2.49) | -2.71 (-2.96, -2.47) |
| | High diversity, NDD | -1.55 (-1.75, -1.36) | -1.46 (-1.66, -1.28) |
| | Low diversity, DI | -1.99 (-2.20, -1.80) | -1.97 (-2.18, -1.76) |
| | Low diversity, NDD | -1.41 (-1.64, -1.19) | -1.07 (-1.32, -0.83) |
| High to low diversity | Large, DI | -1.70 (-1.97, -1.43) | -1.68 (-1.92, -1.44) |
| | Large, NDD | -1.31 (-1.50, -1.11) | -1.57 (-1.76, -1.37) |
| | Small, DI | -0.94 (-1.14, -0.74) | -0.94 (-1.15, -0.73) |
| | Small, NDD | -1.17 (-1.40, -0.95) | -1.17 (-1.43, -0.92) |
| Increasing maladaptation | Large, high diversity, DI | -5.39 (-7.89, -2.91) | -6.66 (-8.86, -4.41) |
| | Large, high diversity, NDD | -5.50 (-7.45, -3.51) | -5.63 (-7.47, -3.81) |
| | Large, low diversity, DI | -6.21 (-8.79, -3.78) | -7.39 (-9.91, -4.95) |
| | Large, low diversity, NDD | -6.18 (-8.46, -3.84) | -6.23 (-8.79, -3.77) |
| | Small, high diversity, DI | -3.79 (-4.66, -2.94) | -3.89 (-4.38, -3.01) |
| | Small, high diversity, NDD | -3.43 (-4.36, -2.49) | -3.39 (-4.38, -2.45) |
| | Small, low diversity, DI | -4.89 (-6.30, -3.52) | -4.01 (-5.45, -2.57) |
| | Small, low diversity, NDD | -4.26 (-5.77, -2.79) | -4.53 (-6.20, -2.82) |

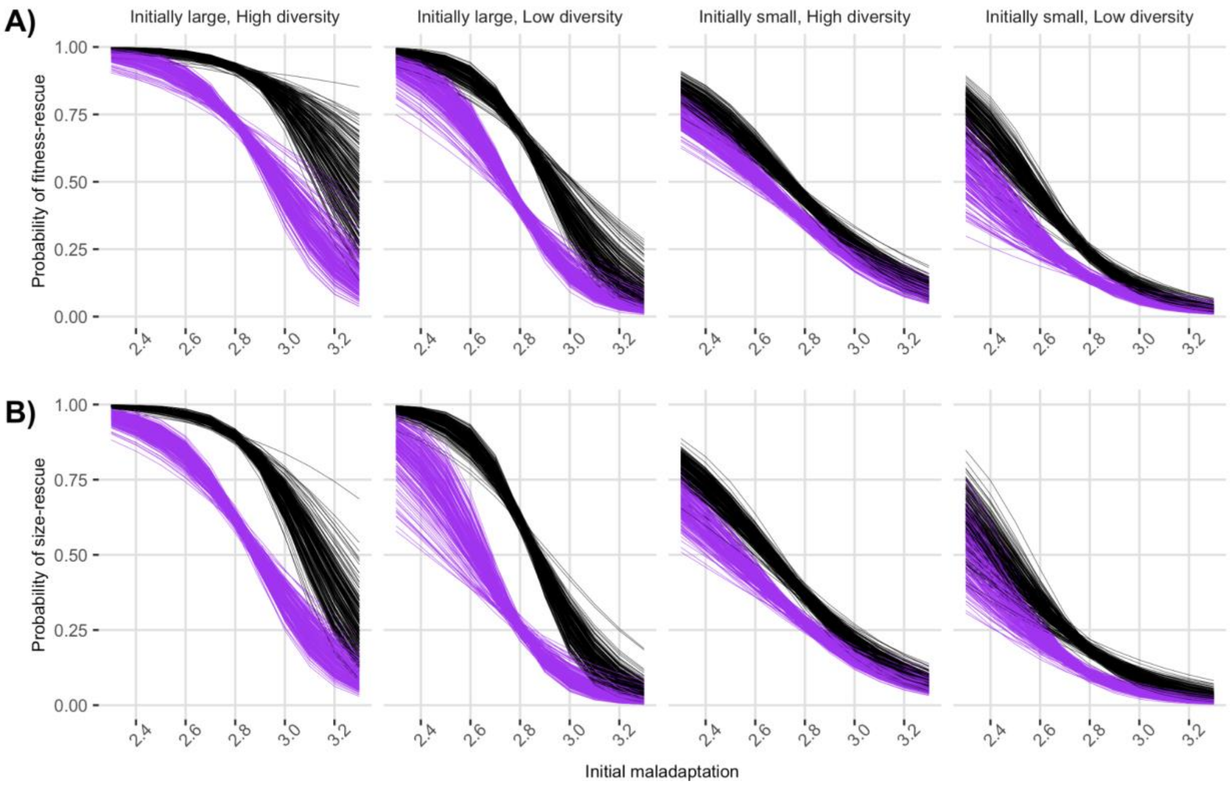


**Figure F1:** Probability of rescue using two different criteria as a function of initial degree of maladaptation, initial size, genetic diversity, and density dependence. Each line corresponds to one posterior estimate from a Bayesian generalized linear model. Black lines correspond to populations with density-independent growth and purple lines correspond to populations with density-dependent growth.

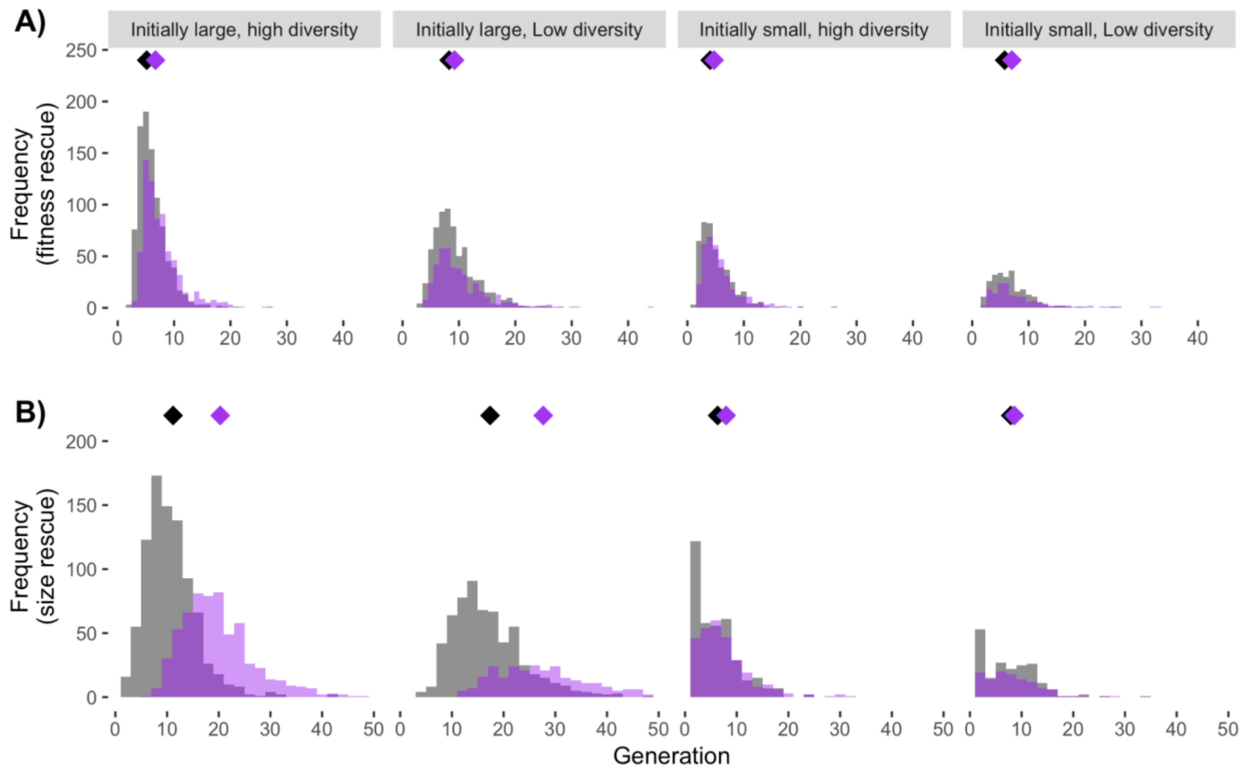


**Figure F2:** Histograms of time until rescue for fitness-based rescue (A) and size-based rescue (B) under density independence (gray) and density dependence (purple). Diamonds indicate the mean rescue time.

### Supporting Information G: State variables conditioned on extinction/survival

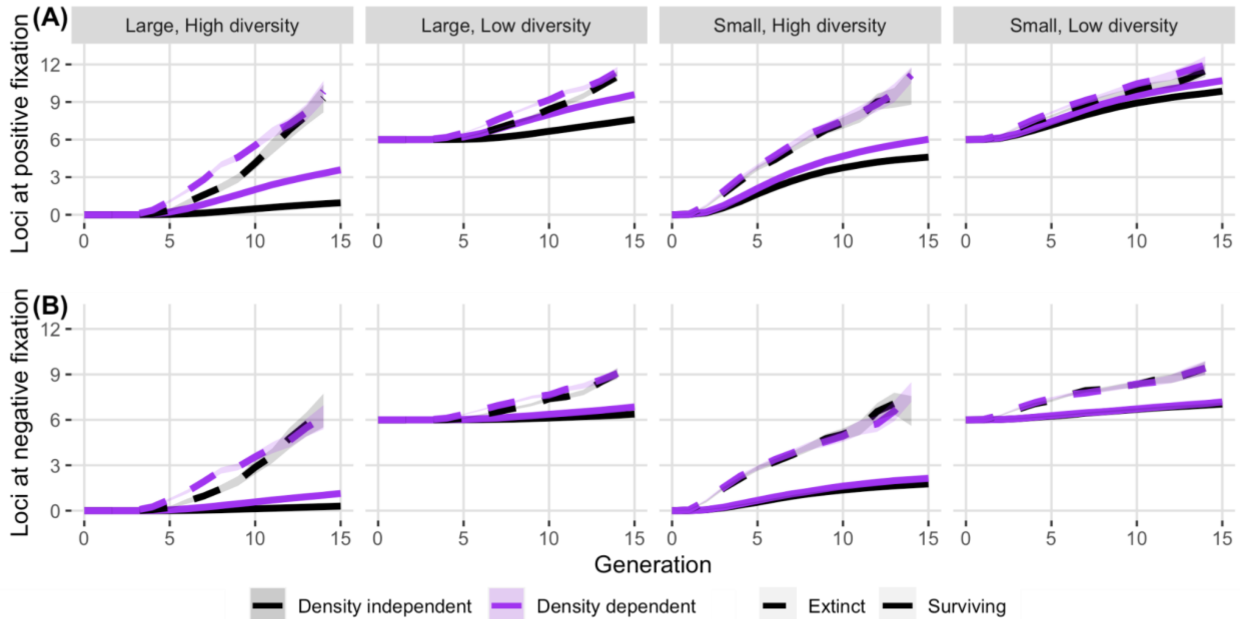


**Figure G1:** Mean number of loci (out of 25) at fixation for the positive (A) and negative (B) alleles, conditioned on extinction or survival to the end of the simulation. Shaded regions represent twice the standard error on each side of the mean.

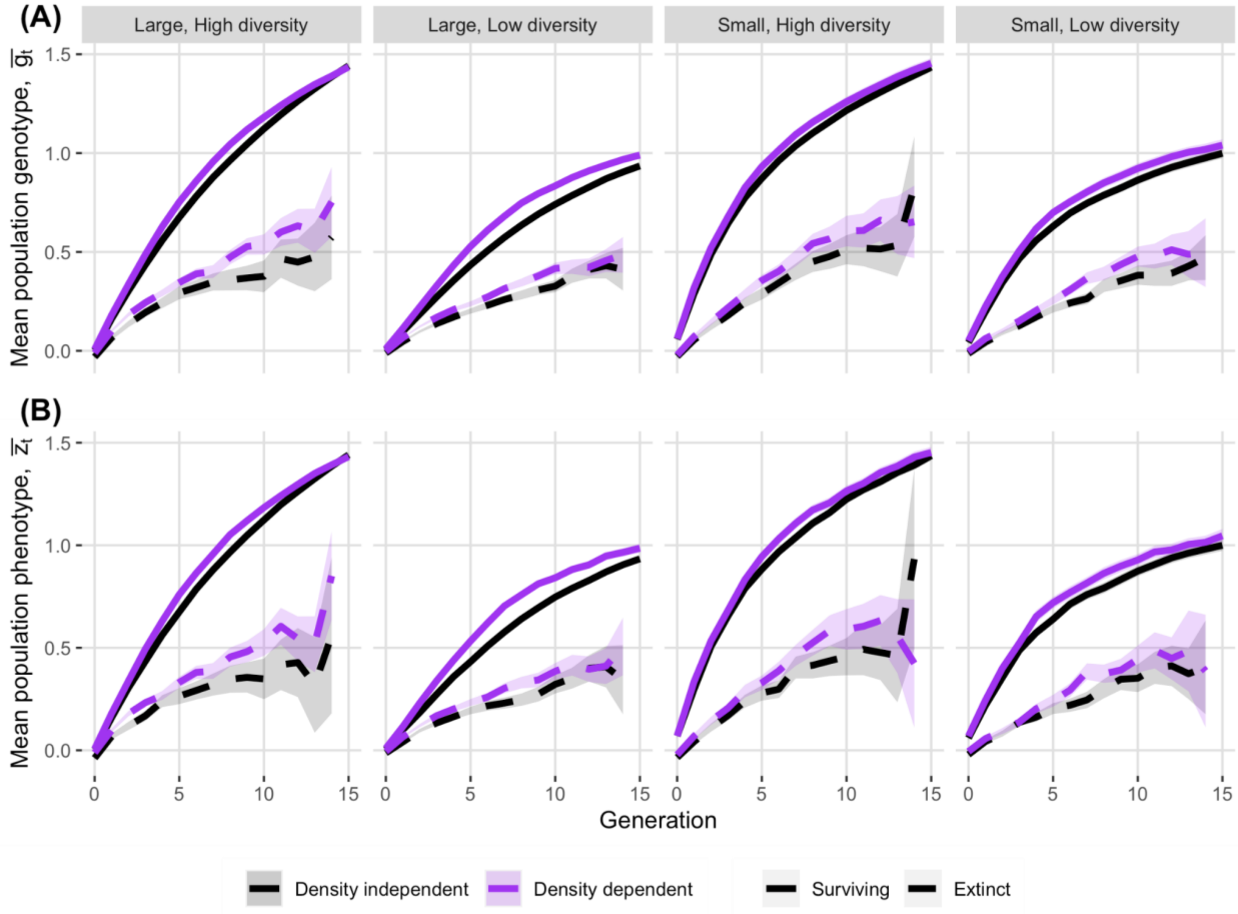


**Figure G2:** Mean population genotype (A) and phenotype (B) over time, conditioned on extinction or survival by the end of the simulation. A positive genotype (phenotype) is closer to the novel environmental optimum, θ = 2.8.

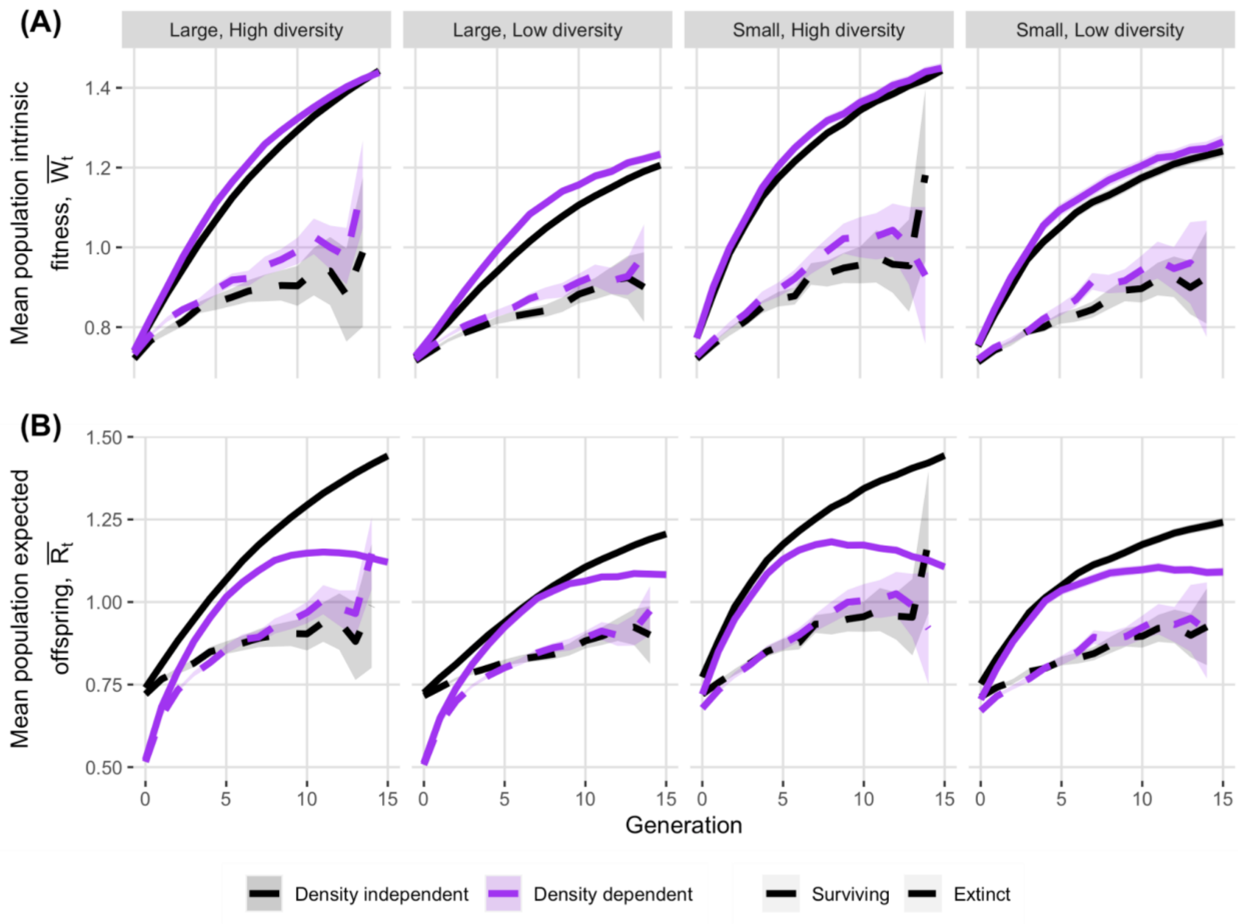


**Figure G3:** (A) Mean population intrinsic fitness, $\underline{W_t}$ and (B) expected number of offspring per individual, $\underline{R_t}$, over time, conditioned on extinction or survival by the end of the simulation.

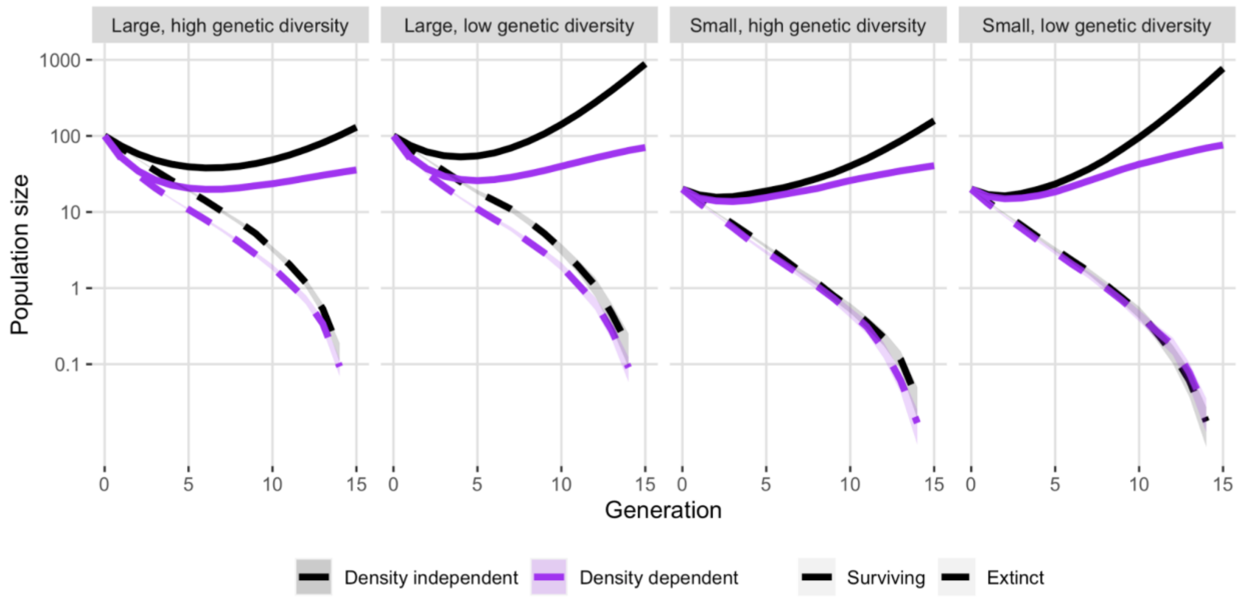


**Figure G4:** Mean population size over time, conditioned on survival to the end of the simulation (solid lines) or extinction before the end of the simulation (dashed lines). Means are estimated with extinct populations included as size zero. Shaded areas represent twice the standard error on either side of the mean; these areas may be difficult to observe in some cases due to low standard error.

### Supporting Information H: State variables conditioned on generation of extinction

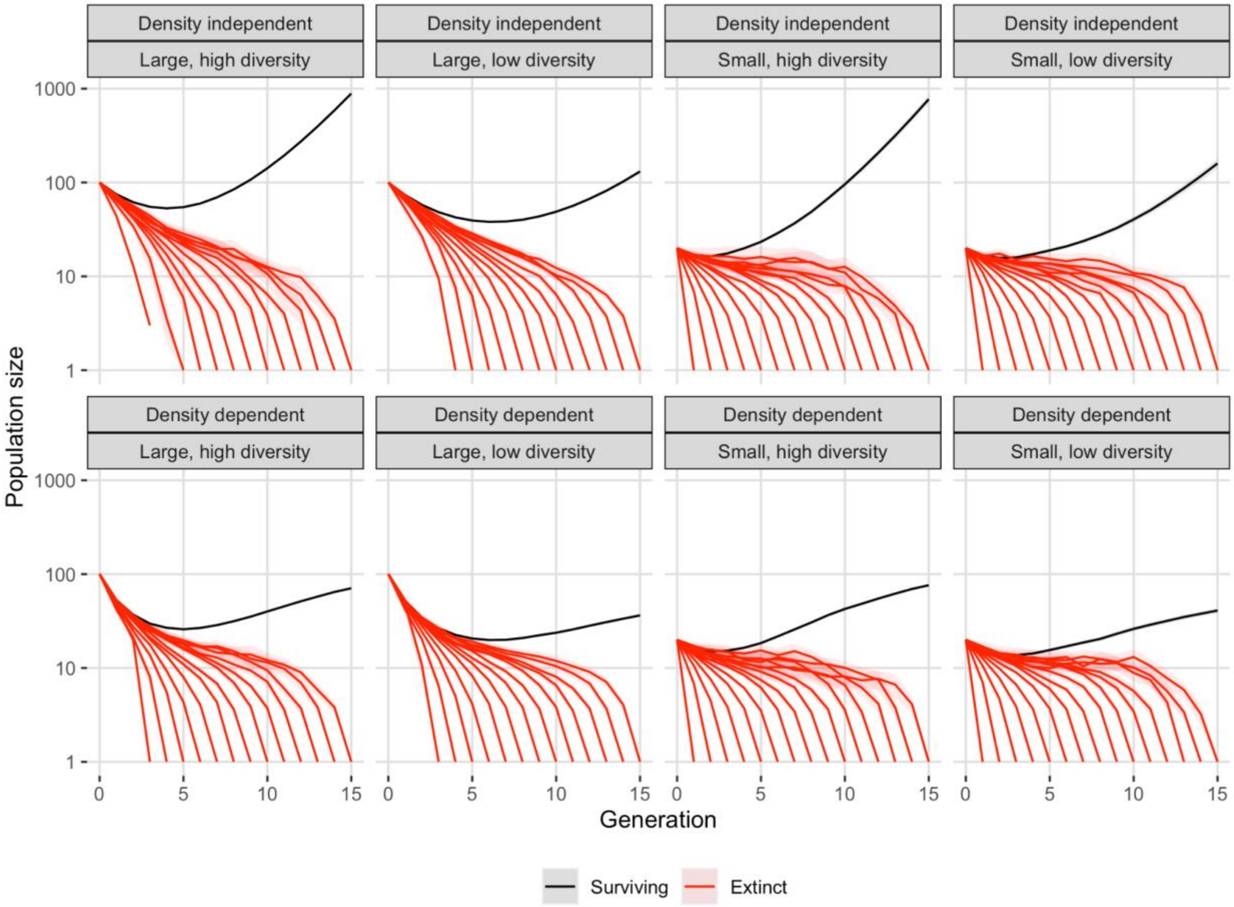


**Figure H1:** Mean population size in each generation, conditioned on generation of extinction for extinct populations (red curves) or survival (black curves). Shaded areas represent twice the standard error above and below the mean.

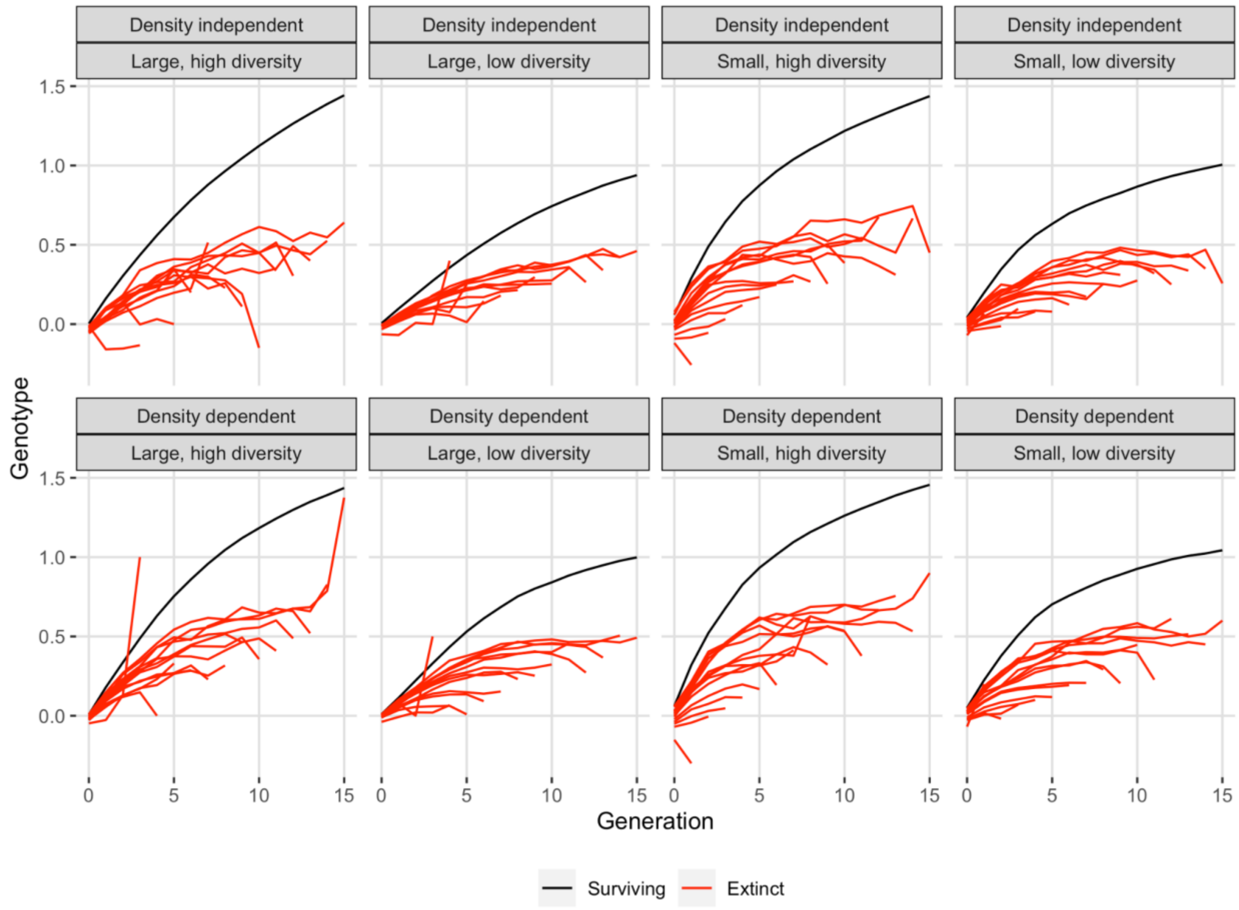


**Figure H2:** Mean population genotype, $\underline{g_t}$, conditioned on survival generation of extinction for extinct populations (red curves) or survival (black curves). Larger values indicate a phenotype closer to the environmental optimum of θ = 2.8.

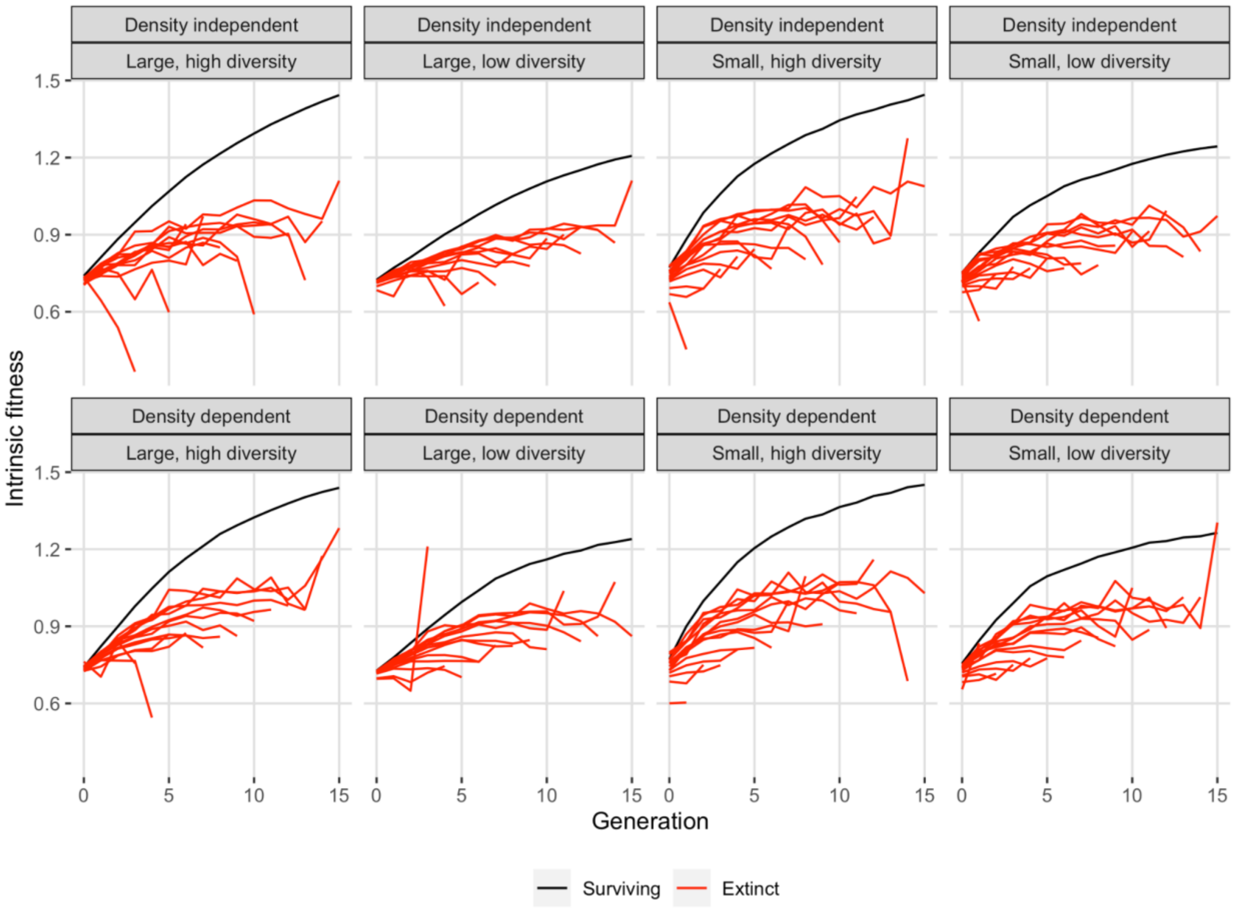


**Figure H3:** Mean population intrinsic fitness, $\underline{W_t}$, conditioned on generation of extinction for extinct populations (red curves) or survival (black curves). In the absence of density dependence, a mean fitness above 1 indicates that the population is expected to grow.

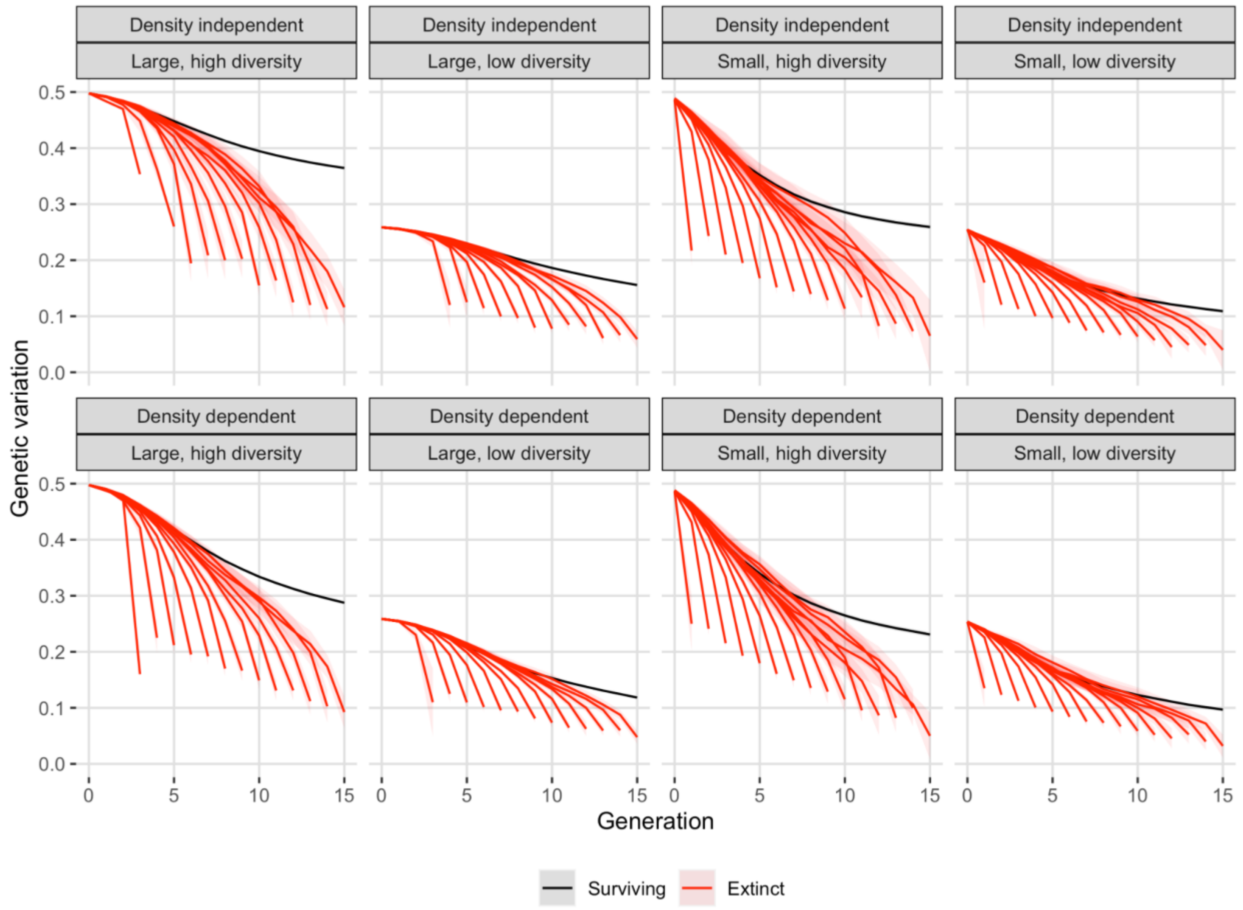


**Figure H4:** Mean population additive genetic variance over time, conditioned on generation of extinction (red curves) for extinct populations or survival (black curves).

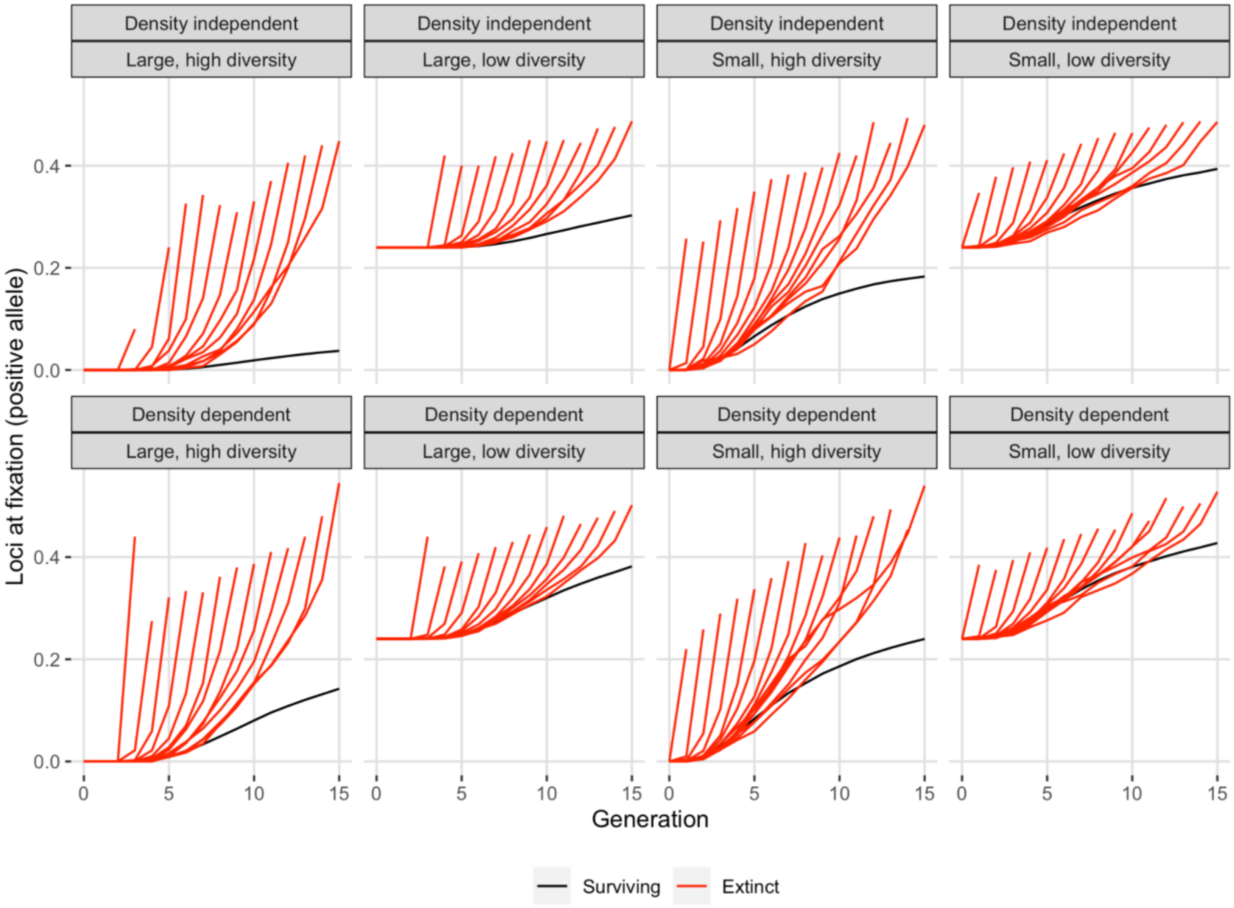


**Figure H5:** Mean proportion of loci (of 25) at fixation for the positive (adaptive allele) over time, conditioned on generation of extinction (red curves) for extinct populations or survival (black curves).

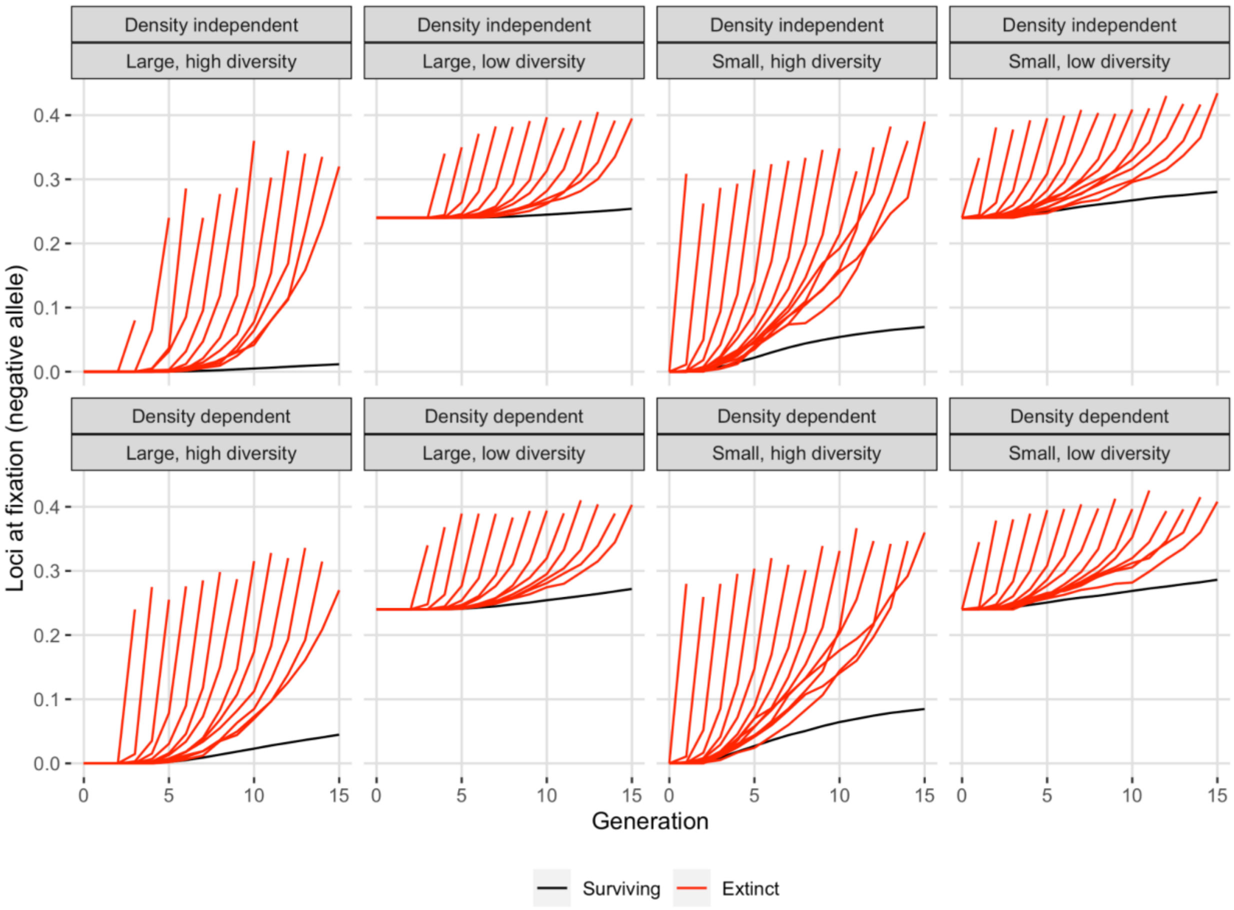


**Figure H6:** Mean proportion of loci (of 25) at fixation for the negative (maladaptive) allele, conditioned on generation of extinction (red curves) for extinct populations or survival (black curves).
